# Redox robustness drives LPMO evolution

**DOI:** 10.1101/2025.06.16.659879

**Authors:** Iván Ayuso-Fernández, Tom Z. Emrich-Mills, Ole Golten, Zarah Forsberg, Kelsi R. Hall, László G. Nagy, Morten Sørlie, Åsmund Kjendseth Røhr, Vincent G. H. Eijsink

**Affiliations:** Faculty of Chemistry, Biotechnology and Food Science, Norwegian University of Life Sciences (NMBU), Ås 1432, Norway; Centre for Biological Research (CIB), CSIC, Madrid 28040, Spain; Biomolecular Interaction Centre and School of Biological Sciences, University of Canterbury, Christ-church 8140, New Zealand; Synthetic and Systems Biology Unit, Institute of Biochemistry, HUN-REN Biological Research Centre Szeged, Szeged 6726, Hungary; Korea University, Seongbuk-gu, Seoul 02481, Republic of Korea

## Abstract

Enzymes known as lytic polysaccharide monooxygenases (LPMOs) are exceptionally powerful small redox enzymes that master the controlled generation and productive use of potentially damaging hydroxyl radicals in what is essentially a H_2_O_2_-driven peroxygenase reaction. We have used ancestral sequence reconstruction and enzyme resurrection to unravel evolutionary steps leading to this unprecedented catalytic power. Real-time monitoring of copper re-oxidation and amino acid radical formation showed evolutionary improvement of both the capacity to avoid futile turnover of H_2_O_2_ and the ability to scavenge damaging radicals resulting from such turnover through a hole hopping pathway. These results show how selective pressure imposed by the need for generating a highly oxidizing intermediate shapes metalloenzymes, involving large parts of the enzyme, well beyond the catalytic center.

## Introduction

The sudden rise in atmospheric O_2_ levels resulting from cyanobacterial photosynthesis during the Great Oxidation Event (GOE), approximately 2.4 billion years ago, has undoubtedly been one of the major drivers of evolution (*1*–*4*). The cataclysmic transition to a new, highly oxidizing atmosphere forced organisms to deal with harmful reactive oxygen species (ROS), potentially damaging all components of the cell (*5*). Although the GOE likely caused a mass extinction (*6*), and while some organisms retreated to anoxic environments, other organisms were better prepared and/or rapidly evolved to deal with oxygen and ROS. The rise in atmospheric O_2_ created a selective pressure in favour of organisms capable of exploiting new, powerful oxygen chemistry, generating the hundreds of reactions performed by oxidoreductases that we observe today in the biochemical networks of living species (*7*).

Today, nature employs oxygen to do some of the most challenging reactions in chemistry, such as the enzymatic oxidation of methane, a perfectly symmetrical molecule, to methanol, which involves activation of a C-H bond with an estimated energy of 105 kcal mol^−1^ (*8, 9*). Another spectacular example concerns the lytic polysaccharide monooxygenases (LPMOs), discovered less than 20 years ago (*10, 11*). Using a single copper co-factor (*12, 13*), these remarkably small interfacial enzymes catalyze the oxidative cleavage of glycosidic bonds in recalcitrant crystalline polysaccharides such as chitin (*11*) and cellulose (*12*–*14*), the two most abundant natural polymers on Earth. This reaction involves activation of a C-H bond with an estimated energy of up to 95 kcal mol^-1^ (*9*) by a highly reactive Cu-oxo species, most likely a predicted but never observed Cu-oxyl (*15*–*18*) (Fig. S1).

Originally, LPMOs were thought to generate the oxidizing copper-oxo species using molecular oxygen (*11*), hence the name monooxygenase, but recent work shows that LPMOs are peroxygenases, using H_2_O_2_ as co-substrate (*19*–*22*). While details of the catalytic mechanism remain unresolved, it is generally accepted that the peroxygenase reaction involves homolytic cleavage of H_2_O_2_ (*16, 18, 19, 23*). Such cleavage leads to the formation of a hydroxyl radical, one of the most powerful radicals emerging in nature, that needs to be contained to avoid damage to the enzyme (Fig. S1) (*16, 19, 22, 24*). The unique interfacial catalytic activity of LPMOs requires a highly exposed catalytic copper site, which makes the enzymes vulnerable to off-pathway redox reactions.

LPMOs are abundant in Nature and it is becoming clear that they are involved in several processes beyond biomass degradation (*25*) including microbial pathogenesis (*26, 27*). In an attempt to understand how these enzymes have evolved to harness the power of copper redox chemistry while preventing damaging off-pathway reactions, we have used ancestral sequence reconstruction (ASR) to interrogate the evolution of chitin-active LPMOs in bacteria. ASR is a powerful computational approach to study the evolution of protein families (*28*). By inferring ancestral sequences in a protein phylogeny, the corresponding enzymes can be “resurrected” in the laboratory to study their properties. Next to providing insight into enzyme evolution, the power of this approach lies in its ability to reveal structural and functional determinants of the efficiency of today’s enzymes.

### Reconstruction of ancient AA10 LPMOs

We first built a phylogeny of 3332 sequences using the whole repertoire of AA10 LPMOs in dbCAN (*29*) (Fig. S2). The dataset was then trimmed to 159 sequences while preserving phylogenetic diversity and including all LPMOs for which functional data were available (Figs. S2 and S3). For ASR, we selected the lineage towards chitin-active LPMOs ending at the extant *Sm*AA10A from the bacterium *Serratia marcescens*, as this LPMO is one of the best characterized to date (*10, 11, 17, 30*). Fig. 1A shows a reduced phylogeny and the four ancestral proteins selected for resurrection: the last common ancestor of all AA10 LPMOs (Anc1), the last common ancestor of chitinolytic AA10 LPMOs (Anc2), the last common ancestor of chitinolytic AA10 LPMOs from Gammaproteobacteria and Bacilli (Anc3) and the most recent ancestor of the same taxonomic diversification (Anc4). Using PAML, and after manual curation of insertions and deletions, we obtained the most probable sequence at each of these nodes (Fig. S4). The deepest node in the phylogeny, Anc1, had by far the highest fraction of ambiguous amino acids, whereas this fraction was smaller and on par with generally accepted levels (*31, 32*) for Anc2-4 (Fig. S5). Before deciding on the final sequences for reconstruction we analysed whether ambiguity affected residues considered crucial for LPMO activity. Apart from Anc1, all such potentially crucial amino acids were reconstructed with a probability close to 1.0 in the ancestral proteins (Table S1-4). Models of Anc1-Anc4, predicted using AlphaFold3 (*33*), all showed similar structural features including a characteristic flat substrate-binding surface that allows binding to the surfaces of crystalline substrates (Fig. 1, B and C). All ancestral LPMOs contain the same catalytically crucial copper binding His-brace (Fig. 1B, Table S1) (*12*) with a Phe in the proximal axial coordination position (Table S4). The number of Cys residues and disulfide bonds changed in evolution (Fig. 1B). Anc1 likely had five Cys residues, forming two disulfide bonds, whereas Anc2 had three disulfide bonds. The number of disulfide bonds was reduced to two in Anc3 and remained at two in Anc4 and *Sm*AA10A. Notably, only the Cys14-Cys22 bond is conserved in all five proteins, and the second disulfide bond in *Sm*AA10A differs from the second in Anc1.

**Figure 1.**
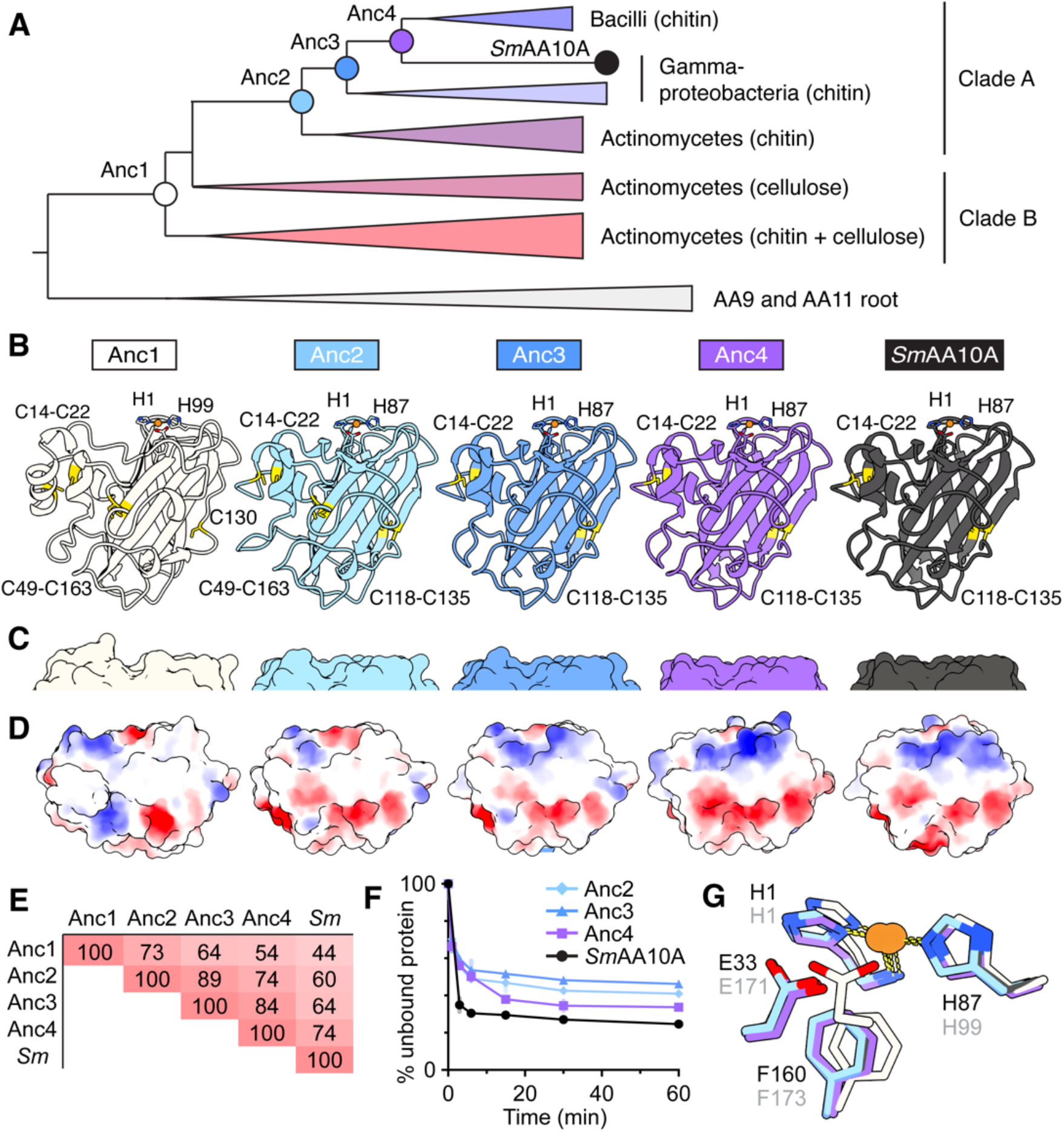
Evolution of *Sm*AA10A and changes to the chitin binding surface. (**A**) Reduced phylogeny schematic for AA10 LPMOs. A full phylogeny is shown in Fig. S3. Clade A corresponds to strictly C1-oxidizing chitin-active LPMOs, while clade B includes LPMOs active only on chitin, only on cellulose, or showing activity on both chitin and cellulose. Functional annotation of clades was based on 25 characterized LPMOs with known activities (*34*). Ancestral nodes are shown as circles. (**B**) Predicted protein models for the four ancestral nodes shown in (A) and the crystal structure of *Sm*AA10A. Copper ions are shown as orange spheres. Flatness (**C**) and surface charges (**D**) of the chitin-binding surface show changes over evolution (red, negative; blue, positive). (**E**) Sequence identity (%) between ancestral LPMOs and *Sm*AA10A. (**F**) Binding of LPMO variants to chitin; error bars are SDs (*N* = 3 technical replicates). (**G**) Overlay of the catalytic centers of *Sm*AA10A and Anc1-4, using the color coding of panel B. Residue numbering in gray corresponds to Anc1 while residue numbering in black corresponds to Anc2-4 and *Sm*AA10A. Note that the position of the catalytically important glutamate in the protein sequence changes in going from Anc1 to Anc2; the location in Anc1 is typical for C1-oxidizing cellulose-active members of the AA10 family. Abbreviations: C, Cys; E, Glu; F, Phe; H, His.

Improvements to surface flatness (Fig. 1C) and changes in binding surface charge (Fig. 1D) are evident over evolution, correlating with an increasing sequence identity of the ancestral proteins to *Sm*AA10A, from Anc1 (44%) to Anc4 (74%) (Fig. 1E, Fig. S4). We detected activity with chitin and no activity with cellulose for all ancestral proteins except Anc1, which showed no activity on any of the tested substrates (Figs. S6-S8). Anc2-4 oxidize chitin in the same manner as *Sm*AA10A, as revealed by product analysis using MALDI-TOF MS (Fig. S6). Binding studies for Anc2-4 showed weaker binding to chitin than *Sm*AA10A, with 46% of Anc3 remaining unbound after 1 hour, compared to 41% for Anc2, 34% for Anc4 and 25% for *Sm*AA10A (Fig. 1F). Variation in binding surface residues likely contributes to the observed differences in substrate affinity between Anc2-4 and *Sm*AA10A (Fig. S9, Table S2, supplementary discussion I).

Redox activity was confirmed for all enzymes, except Anc1, by measuring the oxidase activity, i.e., reductant-driven production of H_2_O_2_ in the absence of substrate (*35*), and the redox potential. Anc2-4 and *Sm*AA10A had similar redox potentials varying from 237 mV to 251 mV, well in line with previous measurements (*36*) (Table S5). They showed oxidase activities in the range of 0.0014 – 0.0122 s^-1^, which compares well with a previously determined rate of 0.001 – 0.002 s^-1^ for *Sm*AA10A (*37*) (Table S5). Taken together, these results show that three of the four ancestral LPMOs are catalytically competent. Comparison of residues in the second sphere of copper coordination (Fig. 1G) shows Anc1 lacks a Glu found at position 33 in *Sm*AA10A and known to be important for catalytic activity (*17*), and instead has an Glu at position 171.

ASR studies frequently show that resurrected enzymes are more thermostable, with a trend showing that deeper nodes are more robust to thermal inactivation (*38*). Measurements of the apparent melting temperature (T_m_) of the resurrected LPMOs and *Sm*AA10A indeed showed the expected trend until Anc3 (Fig. S10). The *apo*-enzymes showed an increase in thermal stability of +12 ºC in Anc4 and +22 ºC in Anc3, compared to *Sm*AA10A. The copper saturated enzymes generally showed higher stabilities, with *Sm*AA10A again having the lowest T_m_. The stability of copper-saturated Anc3 was too high to be assessed with the unfolding assay, and additional assays confirmed the superior thermal stability of this ancestral LPMO (Table S6). For the two most ancestral proteins, however, an increase in apparent T_m_ was not observed. Anc1 and Anc2 showed apparent T_m_ values for the *apo*-enzyme of 49 ºC and 51 ºC, respectively, compared to 63 ºC for apo-*Sm*AA10A. Furthermore, these two ancestral proteins were not stabilized when saturated with Cu^2+^. In the case of the most ancestral protein, Anc1, a destabilizing effect of copper binding (−7 ºC) was observed. ICP-MS analyses showed that Anc1 hardly binds copper. This lack of stability may be taken to suggest flaws in the design of Anc1 and Anc2, although, interestingly, Anc2 is catalytically competent whereas Anc1 is not.

### Evolution of catalytic properties and stability in turnover conditions

In reactions with chitin and an excess of ascorbate as a reductant, *Sm*AA10A releases soluble oxidized products at a constant rate over at least 24 hours (Fig. 2A). In these reactions, LPMO activity is limited by slow *in situ* production of the co-substrate, H_2_O_2_, through abiotic oxidation of ascorbate (see supplementary discussion II). Initially, Anc2-4 were equally capable of converting this available H_2_O_2_ to oxidized chitin, showing that they all are competent LPMOs. However, the ability to sustain the reaction appears to have increased over evolution (Fig. 2A). LPMOs are known to be vulnerable to inactivation resulting from off-pathway reactions with H_2_O_2_ (*19, 24, 39*), so earlier plateauing of the progress curves for Anc2 and Anc3 could reflect a higher vulnerability to inactivation. Faster inactivation may occur because unbound LPMOs react more easily with H_2_O_2_ and/or because such reactions are more damaging (*39*), and/or because of weaker substrate binding (Fig. 1F) (*19*).

**Figure 2.**
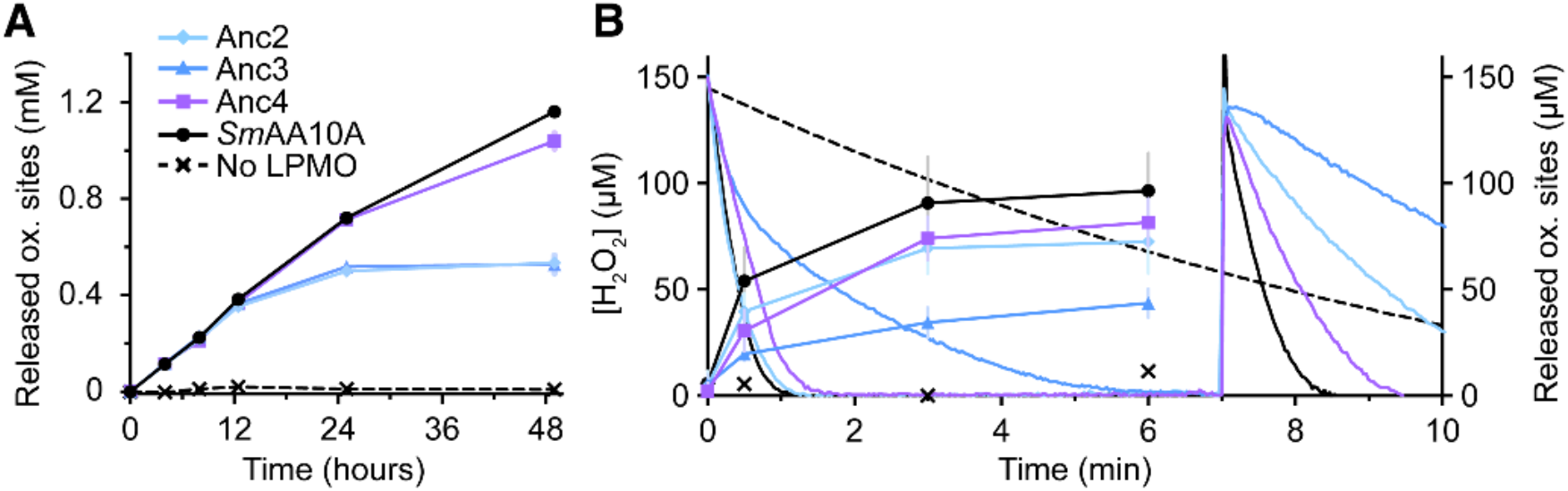
Activity on chitin and stability during turnover. (**A**) Production of soluble oxidized products over 48 hours in reactions without an external source of H_2_O_2_, containing 1 µM LPMO, 10 g L^-1^ chitin and 1 mM ascorbate. Error bars are SDs (*N* = 3 technical replicates). Control reactions without ascorbate did not show significant product formation. (**B**) Dual monitoring of soluble oxidized products (curves with symbols) and real-time H_2_O_2_ consumption (curves only) in reactions with 1 µM LPMO, 10 g L^-1^ chitin, 1 mM ascorbate, 100 mM KCl and 150 µM initial H_2_O_2_ concentration. To address enzyme inactivation, a second addition of 150 µM H_2_O_2_ was made to the reactions after 7 min; the steeper the depletion curve after this addition, the less inactivation has taken place. The dashed black line shows the depletion of H_2_O_2_ in a control reaction without LPMO for the complete experiment (no addition of fresh H_2_O_2_ at 7 minutes) and the black crosses show product levels detected in this control reaction. Data are means and error bars are SDs (*N* ≥ 2; independent experiments). Error bars are not shown for H_2_O_2_ consumption curves for the sake of clarity. The color scheme is as in (A). Reactions were performed in 50 mM sodium (A) or potassium (B) phosphate, pH 7.0, at 37 °C. Abbreviations: SDs, standard deviations.

To clarify these differences, we next measured oxidation of chitin under intentionally damaging conditions, i.e., in the presence of 150 µM H_2_O_2_ and 1 mM ascorbate. Under these conditions LPMOs will catalyze oxidation of the substrate at a much higher rate than in the reductant-driven reactions described above, but, due to the high H_2_O_2_ concentration, oxidative damage of the enzyme will occur. Moreover, under these conditions, the reaction is limited by the amount of active enzyme (as opposed to the amount of available H_2_O_2_ in the reductant-driven reaction), which means that enzyme inactivation will be better reflected in the progress curves. Dual monitoring of H_2_O_2_ consumption and chitin oxidation showed that all enzymes converted the initial 150 µM H_2_O_2_, but with conspicuous differences (Fig. 1B, first six minutes). *Sm*AA10A showed the fastest depletion of H_2_O_2_ and generated the highest product level (i.e., was best at using H_2_O_2_ productively). Anc4 and Anc2 also showed fast, albeit slightly slower, H_2_O_2_ depletion but with reduced product levels, suggestive of a reduced capability to use H_2_O_2_ productively. Of note, the rate of H_2_O_2_ depletion also depends on the rate of the potentially damaging off-pathway peroxidase reaction and, compared to *Sm*AA10A, this reaction is faster for Anc2 and Anc4 (see below). Anc3 performed clearly worse, showing a diminishing H_2_O_2_ depletion rate and low product levels, suggesting non-productive use of H_2_O_2_ causing enzyme inactivation. Addition of a second portion of 150 μM H_2_O_2_ (Fig. 1B, 7 – 10 minutes) showed least inactivation for *Sm*AA10A and Anc4, while Anc2 and Anc3 showed clear signs of inactivation. Residual activity, taken as the estimated initial rate of H_2_O_2_ depletion after the second addition divided by the initial rate at the beginning of the experiment, was 0.14 ± 0.07 for Anc2, 0.15 ± 0.06 for Anc3, 0.61 ± 0.24 for Anc4 and 0.63 ± 0.22 for *Sm*AA10A (means ± SDs).

Taken together, these activity assays with chitin show that the ancestral enzymes are more vulnerable to oxidative damage, and suggest that evolution of efficient chitin oxidation was achieved through an improved ability to use H_2_O_2_ in a productive rather than unproductive fashion.

### Changes in copper reactivity and off-pathway reactions

The observed trend in the rate of copper reduction by ascorbate in the absence of polysaccharide substrate, Anc3<Anc2<Anc4<*Sm*AA10A (Fig. 3A; Table S7), resembles the trends observed for substrate binding (Fig. 1F) and H_2_O_2_ tolerance in reactions with chitin (Fig. 2B). The rates of re-oxidation with H_2_O_2_ in the absence of substrate showed an inversed trend, with such re-oxidation being slowest for the extant enzyme, *Sm*AA10A, and its closest ancestor, Anc4 (Fig. 3B, Table S7). These results support an overarching impression that, as the AA10 LPMOs evolved, their ability to stabilize reduced copper improved and the tendency of the reduced enzyme to engage in potentially damaging reactions with H_2_O_2_ in the absence of substrate, decreased.

**Figure 3.**
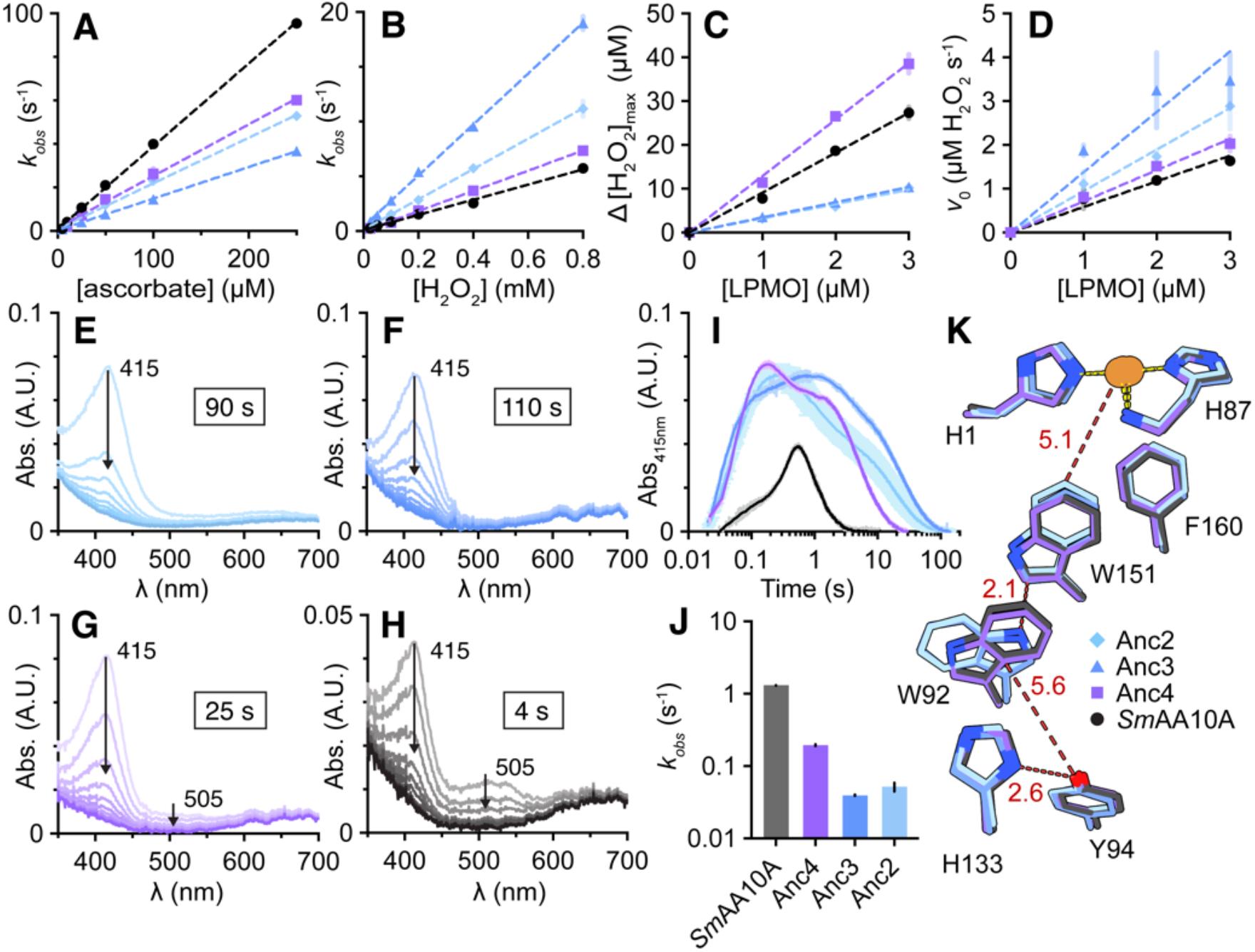
Reduction and re-oxidation kinetics and radical formation in the aromatic protein core in the absence of a polysaccharide substrate. (**A**) Transient state kinetics for reduction of LPMO-Cu^2+^ with ascorbate at 25 °C (*N* = 4). (**B**) Transient state kinetics for oxidation of LPMO-Cu^+^ with H_2_O_2_ at 25 °C (*N* ≥ 3). Dashed lines in (A-B) show linear regression of the data; slopes, defining the apparent second order rate constants, are presented in Table S7. (**C**) Maximum H_2_O_2_ turned over in reactions with 200 µM ascorbate, 100 mM KCl and 100 µM initial H_2_O_2_ without substrate at 30 °C (*N* ≥ 2). Dashed lines show linear regression of the data; slopes define *n*_max_ (listed in Table S8). (**D**) Initial rate of H_2_O_2_ turnover derived from the experiments performed to produce (C). Dashed lines show linear regression of the data; slopes represent the apparent turnover number (TN, s^-1^) (Table S8). Example curves underlying data in (C-D) are shown in Fig. S11. (**E-H**) Absorbance spectra for Anc2 (E), Anc3 (F), Anc4 (G) and *Sm*AA10A (H) after mixing LPMO-Cu^+^ with 20 molar equivalents of H_2_O_2_ at 4 °C in the absence of chitin. The panels show spectra for the decay of the features; spectra for the initial phase of the reactions (formation of the features) are shown in Fig. S12. Panels E-H show ten spectra for each enzyme, spanning the time of maximum absorbance at 415 nm to full decay. The time to full decay is shown in the box in each panel. Traces are means (*N* ≥ 3; technical replicates); errors are not shown for the sake of clarity. Note the different scale of the ordinate axis in (H). (**I**) Time-resolved absorbance at 415 nm derived from the data in panels E-H and Figs. S12-S13. (**J**) The decay rate of the 415 nm absorbance signal extracted from non-linear modelling of the data in (I), illustrated in more detail in Fig. S14. (**K**) Overlay of the aromatic cores of Anc2-4 and *Sm*AA10A from predicted models generated by AlphaFold3 (*33*) and refined using Rosetta Relax (*40*). Distances (in angstroms) between residues are for *Sm*AA10A. Copper ions are shown as orange spheres. Note that Trp92 has the same orientation in Anc2 and Anc3; also note that all shown histidines are close to the protein surface. The color scheme shown in panel K applies to all panels. Error bars in (A-D) and (I-J) are SDs. All reactions were performed in 50 mM sodium phosphate, pH 7.0. Abbreviations: F, Phe; H, His; W, Trp; Y, Tyr; SDs, standard deviations; λ, wavelength; A.U., arbitrary units.

We then used real-time monitoring of H_2_O_2_ consumption to quantify inactivation of LPMOs in the reductant-peroxidase reaction, i.e., re-oxidation of the Cu^+^ active site by H_2_O_2_ in the absence of substrate. By measuring the maximum H_2_O_2_ turned over in reactions with 100 µM H_2_O_2_ (Δ[H_2_O_2_]_max_; Fig. 3C, Fig. S11), we estimated the average number of peroxidase reactions that one LPMO can catalyze before inactivation (*n*_max_) as 9.1 for *Sm*AA10A, in good agreement with the work of Kuusk et al (*39*), who studied the same reaction by monitoring ascorbate depletion. Measurement of the ancestral LPMOs yielded *n*_max_ of 12.9 for Anc4, while Anc2 and Anc3 appeared less resilient to inactivation with *n*_max_ of 3.3 and 3.1, respectively (Fig. 3C, Table S8). Estimation of the initial rates of H_2_O_2_ consumption in these reactions (Fig. 3D) revealed a trend that agreed well with the stopped-flow re-oxidation experiments (Fig. 3B). All in all, while trends varied slightly between experiments and were not perfectly linear over evolution, the data clearly show that Anc2 and Anc3 have the highest re-oxidation rates, i.e., a higher chance of engaging in off pathway reactions, and are more vulnerable to being damaged by such off-pathway reactions.

### Evolution of redox robustness through changes in a hole hopping route

Recent data suggest that bacterial AA10s possess conserved protective hole hopping routes formed by chains of tyrosine and tryptophan residues that traverse the core of the protein, possibly supplemented by histidine residues that facilitate radical dissipation to the solvent through pairing with tyrosine (*30, 41*). During catalysis, highly oxidizing intermediates with a radical nature, referred to as holes, emerge in the enzyme’s catalytic center and are able to extract electrons from aromatic residues in the protein core. By positioning chains of aromatic residues nearby, AA10s seem to have evolved to conduct holes through the protein core to a sacrificial electron donor, avoiding damage to the active site, while generating tyrosyl (Tyr^•^) and tryptophanyl radicals (Trp^•^) along the hole hopping route (Fig. S1). Using stopped-flow UV-vis absorption spectroscopy, short-lived amino-acid radicals can be detected in real time, giving a picture of the kinetics behind the formation and dissipation of these radicals (*23, 30, 42, 43*).

For the extant enzyme, *Sm*AA10A, measurement of the formation (Fig. S12) and dissipation (Fig. 3H) of amino acid radicals in the absence of substrate, revealed a broad Trp^•^ feature around 505 nm accompanied by a stronger Tyr^•^ feature at 415 nm (Fig. 3H). These features align well with previously reported results (*30*). Both features reached maximum intensity in 500 ms (Fig. S12), and their decay took approximately 4 s (Fig. 3, H and I). For Anc2-4 the picture is strikingly different. All ancestors showed a Tyr^•^ feature at approximately twice the absorbance intensity of *Sm*AA10A, which formed slightly faster (appr. 150 ms; Fig. S12) and decayed much slower (Fig. 3E-J), while they showed a diminished or entirely absent Trp^•^ feature (Fig. S13). Anc2 only showed a strong Tyr^•^ feature with a 20-fold longer decay time, disappearing after 90 s (Fig. 3, E and I). Anc3 also showed only the Tyr^•^ peak, with an even longer decay time than its predecessor (Fig. 3, F and I). Anc4 showed a weaker Trp^•^ feature than *Sm*AA10A (Fig. S13), and a Tyr^•^ feature that decayed faster compared to Anc2 and Anc3, (Fig. 3, G and I), though still six times slower than in the extant enzyme. The faster decay of the Tyr^•^ feature over evolution was confirmed by non-linear modelling of the absorbance changes at 415 nm (Fig. 3J, Fig. S14).

All in all, the data show that the kinetics of radical/electron transfer within the aromatic enzyme core have evolved drastically during evolution of these AA10 LPMOs, and that the evolution of higher resistance towards oxidative damage correlates with slower formation and faster dissipation of the Tyr^•^ feature and the eventual emergence of a Trp^•^ feature. Of note, previous studies of *Sm*AA10A have shown that this Tyr^•^ feature is primarily caused by Tyr94 at the exit point of the main hole hopping route (Fig. 3K). Variation in the presence of a Trp^•^ feature between ancestors is remarkable since Trp151, close to the copper, is strictly conserved, and mutational studies have shown that this residue is a major contributor to the Trp^•^ feature in *Sm*AA10A (together with Trp92 (*30*)). Indeed, all LPMOs tested here contain all residues previously identified as constituting a hole hopping route in *Sm*AA10A (*30*), namely Trp151, Trp92, Tyr94 and His133 (Fig. 3K), as well as a third core tryptophan, Trp81 (Fig. S15). The only notable predicted change is a flipping of the side chain of Trp92 when going from Anc3 to Anc4 (Fig. 3K). This change may affect hole hopping since hole transference is sensitive to the spatial configuration of aromatic residues (*44, 45*).

### Substitutions beyond the aromatic core improve redox robustness to near-extant levels

Anticipating that additional residues are involved in shaping radical/electron transfer in the extant enzyme, we identified Phe120 and Asp122 as close neighbours of His133 and Trp92 in *Sm*AA10A, respectively (Fig. 4A; Fig. S15). There is a potential pi-stacking interaction between the benzene ring of Phe120 and the imidazole ring of His133, which could affect the hole-hopping properties of the His133-Tyr94 pair (Fig. S15). Meanwhile, a carboxylic oxygen of Asp122 is within hydrogen-bonding distance (2.9 Å) of the N of the Trp92 indole group (Fig. S15). Interestingly, these two positions appear to have changed drastically and unambiguously through evolution (Table S9). Asp122 appeared for the first time in Anc4, concomitant with the flipping of the Trp92 side chain, while Phe120 appeared for the first time in *Sm*AA10A.

**Figure 4.**
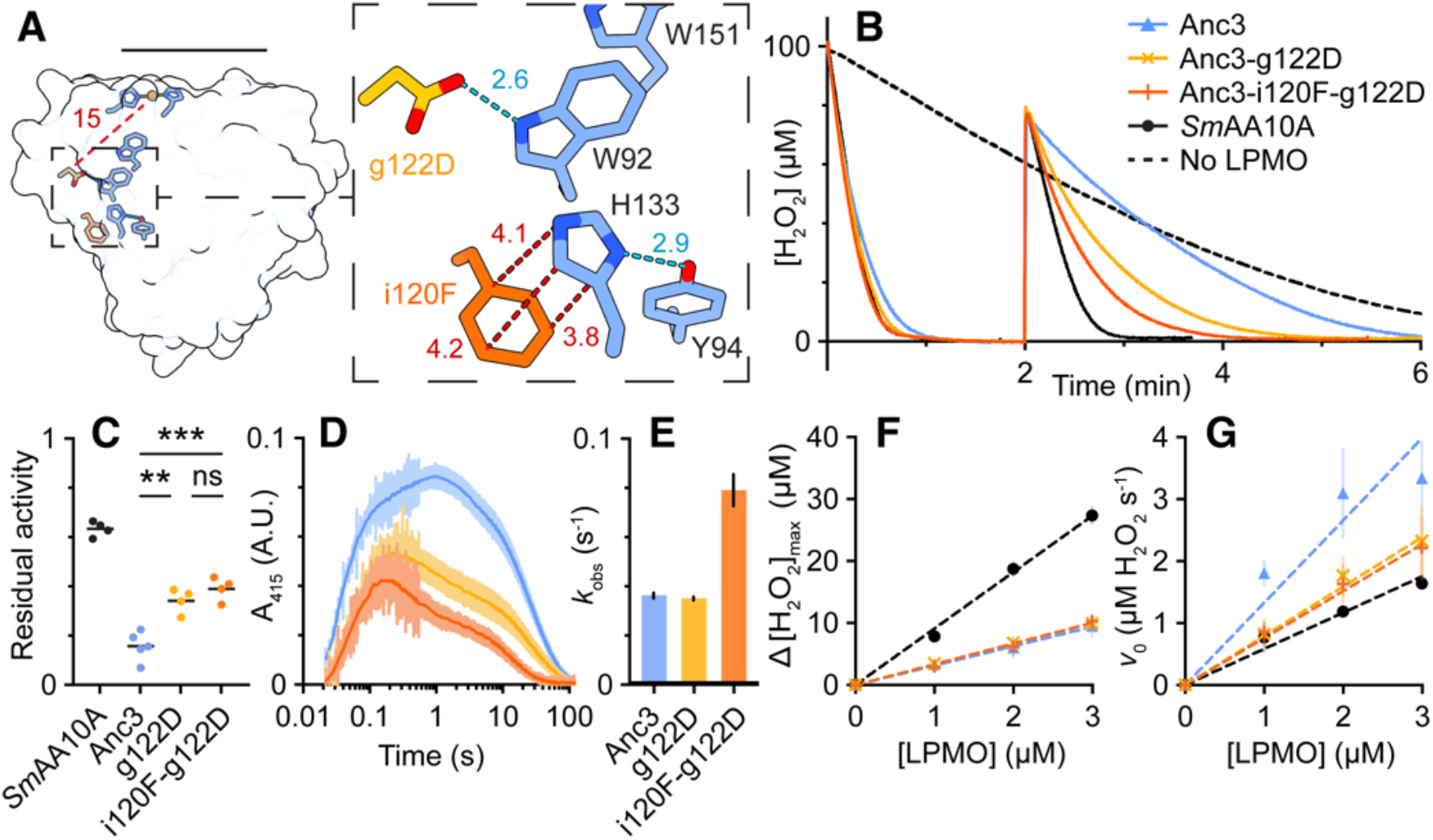
A pair of substitutions that affect the aromatic core and improve redox robustness. (**A**) Predicted model of Anc3 with two substitutions, g122D and i120F, generated by AlphaFold3 (*33*) and refined using Rosetta Relax (*40*). Lower-case letters denote ancestral residue identities, while uppercase letters denote residue identities found in *Sm*AA10A. D122 and F120 are predicted to interact with the aromatic core residues W92 and H133, respectively. Note that the orientation of W92 is flipped relative to non-mutated Anc3. The black line indicates the plane of chitin binding. Distances (red and blue) are in angstroms. Distances in blue correspond to predicted hydrogen bonds (blue dashed lines). (**B**) H_2_O_2_ consumption in reactions containing 1 µM LPMO, 10 g L^-1^ chitin, 100 mM KCl, 1 mM ascorbate and 100 µM initial H_2_O_2_ concentration at 37 °C. A second addition of 100 µM H_2_O_2_ was made after 2 min. Traces are means (*N* = 4; independent experiments); errors are not shown for the sake of clarity. Traces including errors are replicated in Fig. S18. The color scheme applies to all panels. (**C**) Residual activity calculated from the data in (B) as the initial rate of H_2_O_2_ depletion after the second addition of H_2_O_2_ divided by the depletion rate at the beginning of the experiment. Average residual activity (means, shown by horizontal bars) was compared between variants using a two-tailed unpaired *t* test. \*\**P* < 0.01; \*\*\**P* < 0.001; ns, not significant. (**D**) Time-resolved absorbance traces at 415 nm following mixing of LPMO-Cu^+^ with 20 molar equivalents of H_2_O_2_ in the absence of chitin at 4 °C. Error bars are SDs (*N* = 6; technical replicates). (**E**) The decay rate of the 415 nm absorbance signal extracted from non-linear modelling of the data in (D) (see Fig. S19 for details). Bars show means ± SDs. (**F**) Maximum H_2_O_2_ turned over in ascorbate-driven reactions without substrate at 30 °C. Error bars are SDs (*N* ≥ 2; independent experiments). Dashed lines shown linear regression of the data; the slope of each line represents *n*_max_ (listed in Table S8). (**G**) Initial rate of H_2_O_2_ turnover calculated from the same data used to produce panel (F). Dashed lines show linear regression of the data; the slope of each line corresponds to apparent turnover number, (TN, s^-1^) (Table S8). Data for Anc3 and *Sm*AA10A in (F-G) is identical to that shown in Fig. 3, C and D. Reactions were performed in 50 mM potassium (B-C) or sodium (D-F) phosphate, pH 7.0. Abbreviations: g, Gly, H, His; i, Ile; W, Trp; Y, Tyr; A.U., arbitrary units; SDs, standard deviations

Conformational effects of Phe120 and Asp122 on His133 and Trp92 were first assessed through 1 µs Gaussian accelerated MD simulations, considering His133 in its HIE or HID protonation state (i.e. histidine side chain protonated only at the ε-nitrogen or the δ-nitrogen, respectively). The results (Fig. S15) show that in Anc2 and Anc3, Trp92 and each protonation state of His133 are locked in one conformational state. When Asp122 appears in Anc4, Trp92 adopts another single conformational state (cf. “flipping”, mentioned above) that is independent of the protonation state of His133 (Figs. S15 and S16). Structural predictions refined using Rosetta Relax (*40*) for both Anc2 and Anc3 indicated that a mutation to introduce Asp122 induces flipping of Trp92 to the orientation seen in Anc4 (Fig. 4A). The appearance of Phe120 in *Sm*AA10A does not seem to change the conformation of Trp92 or His133 relative to Anc4 (Fig. S15). However, replica-exchange constant-pH simulations (Fig. S17) showed that the presence of Phe120 impacts the p*K*_a_ value (up by one pH unit) and the protonation state of His133 (strong preference for Nε). Simulation of the *in-silico* variants Anc2-v120F, Anc3-i120F and Anc4-i120F (where lower-case letters denote ancestral states and upper-case letters denote extant states) confirmed these observations (Fig. S17).

Next, Anc3 was chosen for mutagenesis due to its markedly low redox robustness compared with *Sm*AA10A, while having a higher probability of being correctly reconstructed than Anc2 (Fig. S5). In reactions with chitin and externally added H_2_O_2_, the Anc3-g122D variant and, even more so, the Anc3-i120F-g122D double mutant showed faster and more continuous H_2_O_2_ depletion than Anc3 (Fig. 4B; first two minutes), indicative of higher stability, as was confirmed by the residual activity after a second addition of co-substrate, which gave rate ratios of 0.16 ± 0.06 for Anc3, 0.34 ± 0.05 for Anc3-g122D, and 0.39 ± 0.05 for Anc3-i120F-g122D (means ± SDs; Fig 4C). Similar effects were observed for the Anc2-g122D mutant, while we could not detect a significant effect on the H_2_O_2_ depletion rate and residual activity of introducing the i120F mutation in Anc4 (Fig. S18). This suggests that the g122D mutation, which does not involve an aromatic residue, but which leads to a structural rearrangement in the aromatic “hole hopping” core of the protein (Fig. 3K) was an important step in the evolution of LPMO redox stability.

Stopped-flow UV-vis absorption spectroscopy overall showed similar hole hopping features for Anc3 and the mutants Anc3-g122D and Anc3-i120F-g122D, but the Tyr^•^ feature showed reduced intensity (Fig. 4D), and its decay was faster in the double mutant (Fig. 4E, Fig. S19). Underpinning the impact of the i120F mutation, the Anc4-i120F mutant, which contains Asp at position 122, showed a clear increase in the decay rate of the Tyr^•^ feature and a much clearer Trp^•^ feature, making the absorbance spectrum strikingly similar to the absorbance spectrum of *Sm*AA10A (Fig. S20). Most remarkably, while the mutations at positions 120 and 122 on the one hand affect redox stability (Fig. 4B) and the putative protective mechanism (hole hopping), all three Anc3 variants showed the same ability to deal with damaging radicals in the peroxidase reaction (same *n*_max_; Fig. 4F, Table S8). Instead of changes in *n*_max_, the improved stability of the Anc3-g122D and Anc3-i120F-g122D variants is due to a reduced tendency to engage in the peroxidase reaction (Fig. 4G, Table S8). This raises questions as to the role of the carefully evolved aromatic core, which undoubtedly plays a role in protective “hole hopping” (*30*), but whose conductivity and redox properties also seem to affect other enzyme features such as the ability of the unbound reduced enzyme to react with H_2_O_2_.

### Concluding remarks

It is well known that the cores of redox enzymes tend to be enriched in aromatic residues (*41*), that participate in radical/electron transport, with a possible protective function (“hole hopping”; (*41*)). LPMOs are vulnerable because they catalyze powerful chemistry and have highly exposed catalytic copper sites and it is thus not surprising that hole hopping phenomena have been observed for these enzymes (*23, 30, 42, 43*). The present study shows that an aromatic core was present early in evolution and that the conductivity of this core has been modulated during evolution. Most importantly, the present study shows that LPMO evolution was almost exclusively driven by redox stability: the initial reaction rates of all catalytically competent reconstructed ancestors are similar to the reaction rate of the extant enzyme (Fig. 2B, 4B), but their redox stabilities vary.

Our results show that during evolution, increased redox stability was achieved by (1) better substrate binding (Fig. 1F), which reduces the chances for the off-pathway peroxidase reaction, (2) an improved ability to avoid damage resulting from the peroxidase reaction (increase in *n*_max_; Fig. 3C), and (3) a reduced tendency to engage in the off-pathway peroxidase reaction in the absence of substrate (Fig. 3, B and D). These changes are accompanied by a clear evolution of the conductivity of the aromatic core of the protein, which is achieved by mutations that modulate the conformation and properties of residues in this core.

It has been suggested that hole hopping is a fundamental mechanism to survive in a richly oxygenated atmosphere (*46*) and our data show that hole hopping changed during evolution. Further studies are needed to fully understand hole hopping in LPMOs and to establish causal relationships between the observed modulations in core conductivity and LPMO redox properties. In this respect, the mutational work with Anc3 is most remarkable: the mutations modulate the formation of radicals in the hole hopping route, but the increase in redox stability is not due to better protection against radicals, but rather a reduced tendency to generate such radicals. This suggests that the aromaticity and fine-tuning of the cores of redox enzymes somehow impact active site reactivity, next to providing radical escape routes.

One open question concerns the true biological function of ancestral LPMOs. It has been proposed that oxidoreductases evolved before the GOE to deal with reactive oxygen species (*47, 48*), and that, later, this powerful chemistry was incorporated into metabolism in myriad ways (*1, 7*). It is perhaps conceivable that mono-copper enzymes performing ROS scavenging initially oxidized other substrates and that side reactions such as chitin oxidation were sequestered and improved later in evolution. An argument for this latter line of thought is the combination of characteristics, displayed by Anc2 and Anc3, of poor ability to sustain chitin oxidation reactions (Fig. 2, A and B) and fast re-oxidation by H_2_O_2_ in the absence of substrate (Fig. 3, B and D). Dealing faster with H_2_O_2_ in the absence of chitin could be pointing precisely to an ancestral role in ROS scavenging.

In conclusion this first detailed experimental characterization of LPMO evolution suggests that LPMOs have evolved to transition from enzymes with low redox robustness and possibly multiple roles centered around dealing with reactive oxygen species, to redox robust peroxygenases for the oxidation of crystalline substrates. LPMOs are very small enzymes, for example compared to another of Nature’s powerhouses, particulate methane monooxygenase (*49*). It is noteworthy that a large part of these small enzymes seems to have evolved for the purpose of redox robustness. Considering that Anc3-i120F-g122D and Anc4-i120F still perform worse than the extant enzyme, additional residues are likely involved, underpinning the notion that exploitation of the catalytic potential of powerful oxygen redox chemistry is accompanied by the evolution of complex protective mechanisms.

## Supporting information

Supplementary Materials (Methods; Supplementary figures, discussions, tables and references)

## Author contributions

I.A.-F., T.Z.E.-M. and V.G.H.E. designed the study. I.A.-F., T.Z.E.-M., O.G., Z.F., and K.R.H. performed the experiments. I.A.-F. and L.G.N. performed the ASR. Å.K.R. performed the MD simulations. All authors interpreted the data. Å.K.R., M.S. and V.G.H.E supervised the work and acquired funding. I.A.-F., T.Z.E.-M. and V.G.H.E. wrote the initial manuscript. All authors contributed to writing the final version of the manuscript.

## Acknowledgements

The authors gratefully acknowledge funding from the European Research Council (ERC) through a Synergy Grant (grant number 856446). We also gratefully acknowledge funding from the Norwegian Research Council for supporting Å.K.R. (project number 301022) and for provision of the stopped-flow instrument via funding from the Norwegian Macromolecular Crystallography Consortium (project number 245828).

