## Supplementary Materials (Methods; Supplementary figures, discussions, tables and references) for "Redox robustness drives LPMO evolution"

*Phylogenetic analysis and ancestral sequence reconstruction*

AA10 LPMO sequences were retrieved from the dbCAN server, that performs an automatic annotation of CAZymes using HMMER and DIAMOND searches (*1*). Initially, a large phylogeny was built containing the catalytic domains of all annotated AA10s (3332 sequences), including enzymes of known activity that were used to predict functional clades as chitin-active, cellulose-active or mixed cellulose-/chitin-active (*2*, *3*). Duplicated sequences and very short or long sequences were removed using cd-hit (*4*). Signal peptides were removed, meaning that all sequences started with His1. Carbohydrate-binding modules (CBMs) were also removed to focus on the evolution of the catalytic domains (CDs) of LPMOs. After this curation, the sequences were first aligned using MAFFT (*5*) after which fasttree (*6*) was used to obtain the phylogenetic tree, both with automatic selection of parameters.

A phylogeny of this size is not suitable to perform ASR since ambiguity in the reconstruction would be too high. To minimize ambiguity, the dataset was reduced to 159 sequences keeping the phylogenetic diversity of the tree and keeping all LPMOs for which functional data were available. The sequences were aligned using MAFFT (L-INS-i option, for accurate alignments) and the resulting multiple sequence alignment (MSA) was used to calculate the evolutionary model that best fits the data, using ProtTest3, according to the corrected Akaike Information Criterion (*7*), resulting in the Whelan & Goldman (WAG) substitution matrix including the gamma distribution (gamma shape 4 rate categories = 1.91) (WAG+G). The phylogeny was built using the MSA and selecting the WAG+G parameters with either PhyML (*8*) or RAxML (*9*), resulting in identical topologies after rooting with AA9 and AA11 LPMOs.

ASR was performed using codeml from PAML4.7 (*10*). The MSA, the rooted phylogeny and the WAG model of evolution were used as inputs to obtain the most probable sequence at each node of the phylogeny and the distribution of amino acids and their posterior probabilities per position in each ancestral sequence. The lineage of the chitin-active LPMO AA10A from *Serratia marcescens*

(*Sm*AA10A, also known as CBP21) was targeted for biochemical characterization. The last common ancestor of all AA10 LPMOs (Ancestral LPMO1, Anc1), the last common ancestor of chitinolytic AA10 LPMOs (Anc2), the last common ancestor of the chitinolytic AA10 LPMOs of Gammaproteobacteria and Bacilli (Anc3) and the most recent ancestor of chitinolytic AA10 LPMOs in a subgroup of the Gammaproteobacteria and Bacilli (Anc4) were selected as representatives in the evolution of chitinolytic AA10 LPMOs. The most probable amino acid sequences at each selected node were corrected for insertions and deletions artificially introduced in the ASR. The most probable sequences were obtained after analysis of posterior probability distributions using the output matrix from codeml. Ambiguity at key positions was analysed based on the distribution matrix, where the probabilities of each amino acid at each position in every node are shown. The selection of key positions was based on a literature search for amino acids involved in LPMO reactivity and is described in detail in tables S1-S4. For these key positions, a probability cutoff of > 0.995 was considered unambiguous. The resulting most probable sequences (provided in Fig. S4) were used to design genes optimized for expression in *Escherichia coli*.

*Protein expression and purification*

With the exception of Anc1 (see below), all ancestral proteins and *Sm*AA10A were cloned, expressed and purified as described previously (*11*). In short, gene fragments with the sequence of codon optimized genes for *E. coli* expression were synthesized by Twist Biosciences and cloned into the pRSET-B backbone using a Gibson Assembly® kit (New England Biolabs, Ipswich, MA, USA). The successfully cloned and sequence-verified constructs were transformed into One Shot™ BL21 Star™(DE3) chemically competent *E. coli* cells (Invitrogen, Waltham, MA, USA), which were used as expression hosts. The expression protocol started using a single colony to inoculate 1.8 L baffled shake flasks containing 1 L of LB medium supplied with 100 µg $mL^{-1}$ of ampicillin. The expression of LPMOs in pRSET-B is constitutive, and after inoculation the culture was grown for 16 hours at 37 ºC, 200 rpm. The cells were harvested, and each protein was collected by periplasmic extraction with cold osmotic shock as previously described (*11*). The proteins expressed this way were purified in one single step after loading the periplasmic extracts, supplemented with $(NH_4)_2SO_4$ to a final concentration of 1 M (*Sm*AA10A) or 1.5 M (ancestral proteins), onto a 10 mL chitin resin column (New England Biolabs) equilibrated with 50 mM Tris, pH 8.0, containing 1.0 M or 1.5 M $(NH_4)_2SO_4$. Unbound protein was removed by extensive washing with the same buffer, and pure protein was eluted with 20 mM acetic acid. Eluates were immediately dialyzed vs 50 mM Tris, pH 8.0. Copper saturation was achieved by incubation with a 3-fold molar excess of $CuSO_4$ for 30 minutes at room temperature, after which size exclusion chromatography was performed using a HiLoad 16/600 Superdex 75 column (Cytiva, Marlborough, MA, USA) to remove excess copper and exchange into 50 mM sodium phosphate buffer, pH 7.0. Purified proteins were stored at 4 °C.

Anc1 was expressed using a recombinant protein expression service (GenScript, Piscataway, NJ, USA). A His-SUMO tag was fused to Anc1 immediately before the His1 in the pSUMO plasmid and *E.coli* BL21 Star™ (DE3) competent cells (Invitrogen) were transformed with the recombinant plasmid. A single colony was inoculated into LB medium containing 50 μg mL$^{-1}$ kanamycin and cultures were incubated at 37 °C and 200 rpm. Once cell density reached $OD_{600}$ = 0.6-0.8, 0.5 mM IPTG was added for induction. A Ni column was used to purify the fused His-SUMO-Anc1, followed by tag removal with a SUMO protease, with a second Ni column purification step to obtain the purified Anc1 as unbound fraction. Purified Anc1 was provided by GenScript as a 1 mg mL$^{-1}$ stock solution in 50 mM Tris, pH 8.0, with 5% glycerol.

Protein concentrations were quantified by measuring the absorbance at 280 nm, using theoretical extinction coefficients of 41,160, 42,315, 42,190, 36,690 and 35,200 M$^{-1}$ cm$^{-1}$ for Anc1, Anc2, Anc3, Anc4 and *Sm*AA10A, respectively.

*Binding to chitin*

Binding assays were performed in reactions containing 10 g L$^{-1}$ chitin in 50 mM sodium phosphate, pH 7.0, at 22 ºC and 800 rpm. The binding reactions were initiated by addition of 5 μM LPMO to the mixture ($t_0$), after which the reaction was sampled at several time points over the course of 60 minutes. Samples were subjected to filtration through a MultiScreen™ 96-well filter plate (Sigma-Aldrich, St. Louis, MO, USA) after which protein concentrations in the filtrates, representing unbound protein, were determined with and adapted Bradford protocol for measuring low protein concentrations (*12*). For each protein, a control reaction without chitin was used as reference to calculate the fraction (%) of unbound protein. Controls without enzyme were also added to monitor unspecific signals in the Bradford assay. All reactions were performed in triplicates.

*Initial detection of activity and product analysis by MALDI-TOF*

To detect chitin oxidation, 1 μM of pure enzyme was pre-incubated for 30 min with 10 g L$^{-1}$ chitin in 50 mM sodium phosphate buffer, pH 7.0, at 37 ºC and 800 rpm agitation, after which the LPMO reaction was initiated by addition of 1 mM ascorbate. Reactions were incubated for 24 hours at 37 ºC and 800 rpm agitation. Samples taken from the reaction were filtered through a MultiScreen™ 96-well filter plate (Sigma-Aldrich, St. Louis, MO, USA). In the case of Anc1, where activity on cellulose or chitin could be expected, the same setup was used to test activity on 10 g L$^{-1}$ of Avicel. Soluble products resulting from degradation of Avicel or chitin were analysed using a matrix-assisted laser desorption/ionization time-of-flight (MALDI-TOF) UltrafleXtreme mass spectrometer (Bruker Daltonics, Billerica, MA, USA). 1 μL of the filtered reaction sample was mixed with 2 μL of a matrix solution [9 mg mL$^{-1}$ 2,5-dihydrooxybenzoic acid containing 1 mM NaCl in 30% (v/v) acetonitrile] on the surface of an MTP 384-ground steel target plate (Bruker Daltonics). The target plate was air-dried, and MS data were collected using Bruker flexControl software as described previously (*11*).

*Cellulose and chitin activity assays for Anc1*

Activity of Anc1 loaded with copper was further assessed in reactions with 0.5% (w/v) α- and β-chitin, 0.2% (w/v) bacterial microcrystalline cellulose (BMCC), 0.5% (w/v) filter paper (Whatman no.1), 1% (w/v) Avicel® PH-101 purchased from Sigma Aldrich (St. Louis, MO, USA), or 0.2% (w/v) phosphoric acid swollen cellulose (PASC), which was prepared from Avicel essentially as described in (*13*). Reactions were performed with 1 µM Anc1 or, as positive controls, *Sm*AA10A (chitin) and *Sc*AA10C (cellulose; (*14*)) in 50 mM Tris, pH 8.0, and 5% glycerol, in the presence of 1 mM ascorbic acid [dissolved in TraceSELECT™ water (Honeywell Riedel-de-Haën, Charlotte, NC, USA)] and incubated for 48 hours in an Eppendorf Thermomixer (Eppendorf, Hamburg, Germany) set to 30 °C and 800 rpm. Samples were filtrated using a 96-well filter plate (Sigma-Aldrich) operated with a Millipore vacuum manifold to remove insoluble substrate and the supernatant was analyzed for oxidative products using HPAEC-PAD (see below for details).

*Product analysis*

Oxidized chito- and cello-oligosaccharides were analyzed with a high-performance anion-exchange chromatography method (*15*) using a Dionex™ ICS-5000 system (ThermoFisher Scientific, Waltham, MA, USA) set up with a disposable electrochemical gold electrode. Samples of 5 µL were injected on a CarboPac PA200 (3×250 mm) column operated with 0.1 M NaOH (eluent A) at a flow rate of 0.5 mL $min^{-1}$ and a column temperature of 30 °C. Elution was achieved using a stepwise gradient with increasing amounts of eluent B (0.1 M NaOH + 1 M NaOAc), as follows: 0–10% B over 10 min; 10–30% B over 25 min; 30–100% B over 5 min; 100–0% B over 1 min; and 0% B (reconditioning) for 9 min. Chromatograms were recorded using Chromeleon 7.0 software (ThermoFisher Scientific). Standards of C1-oxidized chito-oligosaccharides (DP2, 4 and 6) were generated *in-house* by complete oxidation of *N*-acetyl-chito-biose, -tetraose, or -hexose (Megazyme; > 85-95% purity) with the *Fusarium graminearum* chitooligosaccharide oxidase (*Fg*ChitO) (*16*), as described previously (*17*). Cellulosic LPMO products were assessed using standard mixtures of C1-oxidized $Glc_2$-$Glc_6$, which were produced *in-house* using *Myriococcum thermophilum* cellobiose dehydrogenase (*18*), according to a previously published protocol (*19*).

*Inductively coupled plasma mass spectrometry (ICP-MS)*

ICP-MS was utilized to determine the copper content in Anc1. 2 µM of copper saturated Anc1 in TraceSELECT™ water (Honeywell Riedel-de-Haën) was analyzed using a tandem quadrupole 8800 ICP-QQQ instrument (Agilent Technologies, Santa Clara, CA, USA), equipped with a collision/reaction cell, operated in triple quadrupole mode with helium as collision gas to minimize diatomic interferences from plasma or sample. A multielement internal ICP-MS standard (Inorganic Ventures, Christiansburg, VA, USA) and 65% NORMAPUR® nitric acid (VWR, Radnor, PA, USA) were mixed with the samples before autoclaving at 121 °C for 30 min in sealed tubes. After cooling, samples were diluted with deionized 18.2 MΩ type I water to 10% (v/v) nitric acid. To check and compensate for instrumental drift, a control standard (Inorganic Ventures) was analyzed between samples. Calibration curves were prepared prior to sample analysis, and copper concentrations were determined according to these curves using indium as an internal standard. The experiment was performed in duplicate *(N* = 2).

*2,6-DMP oxidation and thermal inactivation assays*

2,6-DMP oxidation was used to assess thermal stability (Table S6). 2,6-DMP is a small phenolic molecule that LPMOs can oxidize to generate the colored product coerulignone in two consecutive reactions consuming $H_2O_2$ as co-substrate (*20*). The oxidation of 2,6-DMP was monitored by following the formation of coerulignone at 473 nm according to a previously published protocol (*20*). Activity toward 2,6-DMP was used to evaluate the thermal stability of the ancestral LPMO Anc3 vs *Sm*AA10A, using 100 µM $H_2O_2$. Prior to measuring their activity, the proteins, 2 µM in 50 mM sodium phosphate, pH 7.0, were incubated either at 75 or 90 ºC for 10 mins, followed by 1 minute centrifugation at 14000 rpm and cooling down to RT. Reactions for determination of remaining activity contained 2 mM 2,6-DMP in 50 mM sodium phosphate, pH 7.0, with 100 µM $H_2O_2$ and were pre-incubated for 5 minutes at 30 ºC, after which the reactions were initiated by addition of 2 µM LPMO and formation of coerulignone was followed at 473 nm over 2 minutes. Velocities were calculated for the initial, linear phase of the reaction as $\Delta Abs_{473nm}$ $min^{-1}$. All reactions were performed in triplicate.

*Oxidase activity assay*

The ascorbate-oxidase reaction (reductant-driven production of $H_2O_2$) was monitored using the Amplex Red assay (*21*). Reactions contained 2 µM LPMO, 100 µM Amplex™ Red Reagent, 5 U $mL^{-1}$ horseradish peroxidase (Sigma-Aldrich, St. Louis, MO, USA) in 50 mM Tris, pH 8.0, at 30 ºC. Reactions were initiated by addition of 1 mM ascorbate. Formation of resorufin was monitored at 540 nm over 20 min in a Multiskan™ FC microplate photometer (ThermoFisher Scientific, Waltham, MA, USA) and calibrated using a standard curve of $H_2O_2$ that also contained 1 mM ascorbate. Reaction rates are presented in Table S5, corrected for the rates in reactions with no LPMO.

*Determination of the redox potential*

The cell potential of the LPMO-$Cu^{2+}$/LPMO-$Cu^{+}$ redox couple was determined from the reaction between reduced *N,N,N′,N′*-tetramethyl-1,4-phenylenediamine ($TMP_{red}$) and LPMO-$Cu^{2+}$. All stock solutions were deoxygenated by $N_2$ sparging using a Schlenk line and subsequently prepared to working dilutions in an anaerobic chamber. Reactions contained 35 µM LPMO and 150 µM $TMP_{red}$ in 20 mM Tris buffer, pH 8.0, and were incubated at room temperature for 5 min. The generated $TMP_{ox}$ was measured anaerobically using a Nanophotometer C40 (Implen, München, Germany) at 610 nm. The cell potential of the LPMO-$Cu^{2+}$/LPMO-$Cu^{+}$ redox couple was determined from the concentration of $TMP_{ox}$, as described previously (*14*, *22*). The reported values are referenced against a normal hydrogen electrode (NHE).

*Thermal stability*

SYPRO® orange dye from Invitrogen was used to monitor the apparent melting temperature ($T_m$) of LPMOs (*23*). Fluorescence by this dye is naturally quenched and is enhanced when it binds to hydrophobic residues of proteins that become exposed upon unfolding. A temperature ramp from 25 to 98 ºC with an increase of 1.5 ºC $min^{-1}$ was used to evaluate the $T_m$. The SYPRO® orange stock solution (5000x) was diluted to an 8x stock in Milli-Q $H_2O$ prior to adding it to the reaction to reach a 1x working concentration. The reactions contained 30 µM $Cu^{2+}$-saturated LPMO and 1x SYPRO® Orange dye in 50 mM Tris, pH 7.0. Reactions were prepared in quadruplicates, including control reactions without enzyme. The fluorescence change was monitored with a StepOnePlus™ Real-Time PCR (ThermoFisher Scientific, Waltham, MA, USA). The StepOnePlus™ software was used to obtain the negative first derivative of the normalized fluorescence signal with respect to temperature (*-dF/dT*), allowing easy identification of the apparent $T_m$ as a *-dF/dT* peak.

*Time-course reactions of chitin oxidation and quantification of oxidized products*

In LPMO reactions under apparent monooxygenase conditions, the reaction is reductant-driven and there is no external source of $H_2O_2$. These reactions are limited by *in situ* generation of $H_2O_2$ due to oxidation of the reductant. Such reactions were carried out by incubation of 1 µM LPMO with 10 g $L^{-1}$ chitin from squid pen (Batch 20140101, France Chitin, Orange, France), milled with zirconium oxide grinding tools in a PM200 planetary ball mill (Retsch, Haan, Germany) to a particle size of 75 µm, in 50 mM sodium phosphate buffer, pH 7.0. After incubation for 30 mins at 37 ºC and 800 rpm, to ensure substrate-binding, the reaction was initiated by addition of 1 mM ascorbate. The reactions (in triplicates) were sampled at different time-points over 48 hours. Reactions with a high amount (150 µM) of added $H_2O_2$ were done to create damaging conditions. These reactions are not $H_2O_2$-limited and are much faster. Following 1 hour of pre-incubation and addition of 1 mM ascorbate to initiate the reaction, the reactions were sampled over six minutes. For details of the setup, see the section on real-time $H_2O_2$ consumption, below. In both setups, enzyme reactions were stopped by filtration through a MultiScreen™ 96-well filter plate (Merck) that separates the enzyme and soluble reaction products from the remaining substrate. The soluble products present in the filtered samples were degraded by incubation with 1 µM chitobiase from *Serratia marcescens* (*Sm*CHB) overnight at 37 ºC (*17*). *Sm*CHB degrades soluble oxidized chito-oligomers to GlcNAc (native monomer) and GlcNAcGlcNAc1A (chitobionic acid, the oxidized dimer). Analysis of chitobionic acid was performed using a 100 × 7.8 mm Rezex RFQ-Fast Acid H+ (8%) (Phenomenex, Torrance, CA, USA) column operated at 85 ºC in an RSLC system (Dionex, Sunnyvale, CA, USA), with isocratic elution with 5 mM sulphuric acid. 8-

μL samples were injected and analytes were eluted in six-minute runs, with a 1 mL min$^{-1}$ flow rate. The analytes were monitored using a 194 nm UV detector. A standard curve (25–800 μM) of chitobionic acid was included to quantify analytes. The standard was generated by incubating *N*-acetyl-chitobiose (95% purity, Megazyme, Wicklow, Ireland) with a chito-oligosaccharide oxidase from *Fusarium graminearium* for complete oxidation, as previously described (*17*, *24*).

*Real-time $H_2O_2$ consumption measurements*

$H_2O_2$-driven LPMO activity in the presence or absence of substrate was monitored by fast electrochemical detection of the depletion of $H_2O_2$ using an Autolab PGSTAT101 potentiostat (Metrohm, Herisau, Switzerland) as originally described by Schwaiger *et al* (*25*). The setup and procedures used were similar to those detailed in Ayuso-Fernández, *et al* (*26*). Briefly, measurements were performed in a 4 mL volume at constant temperature using a three-electrode setup: a gold rotating disk electrode (with an angular velocity of 50 s$^{-1}$) as the working electrode, a platinum sheet as the counter electrode and a Ag|AgCl electrode in 3 M KCl as the reference electrode. The working electrode was coated in a layer of Prussian blue and protected using a layer of Nafion (Sigma-Aldrich, St. Louis, MO, USA). All reactions were performed in 50 mM potassium phosphate, pH 7.0, and 100 mM KCl as the electrolyte. A low concentration of EDTA (5 μM), that does not deplete copper from active LPMOs on these timescales, was included in all reactions to scavenge free copper released from inactivated LPMOs. Data were acquired using NOVA 1.1 software from Metrohm and processed using custom scripts to correct for system drift and convert data from amps to $[H_2O_2]$ using a standard curve recorded at the beginning of each run.

Reactions in the absence of chitin were inspired by previously described measurements of ascorbate-peroxidase activity by Kuusk et al (*27*). These contained 100 μM initial $H_2O_2$ and 0, 1, 2 or 3 μM LPMO, and were initiated with 200 μM ascorbate at 30 °C. Non-linear modelling of $H_2O_2$ depletion was performed according to the equation $[H_2O_2] = \Delta[H_2O_2]_{max} \cdot e^{-kobs \cdot t} + [H_2O_2]_{\infty}$, where $\Delta[H_2O_2]_{max}$ is the maximum $H_2O_2$ turned over in each reaction and $[H_2O_2]_{\infty}$ is the remaining $H_2O_2$ concentration. The initial rate, $v_0$, of $H_2O_2$ depletion (in μM $H_2O_2$ s$^{-1}$) was calculated as $\Delta[H_2O_2]_{max} \cdot k_{obs}$. Plots of $\Delta[H_2O_2]_{max}$ vs [LPMO] and $v_0$ vs [LPMO] were fitted by linear regression using the least squares method. The resulting slopes are presented in Table S8; the slope of $\Delta[H_2O_2]_{max}$ vs [LPMO] defines $n_{max}$, the average number of peroxidase reactions that one LPMO can catalyze before inactivation, while the slope of $v_0$ vs [LPMO] defines the turnover number (TN, s$^{-1}$).

Reactions in the presence of chitin contained 100 or 150 μM initial $H_2O_2$, 0 or 1 μM LPMO and 10 g L$^{-1}$ chitin and were initiated with the addition of 1 mM ascorbate at 37 °C. LPMOs were incubated with chitin for at least 1 hour at 22 °C with agitation prior to measurement to allow for maximum binding. A second addition of $H_2O_2$ was made after the initially added $H_2O_2$ was fully depleted. Quantification of residual activity after the first reaction cycle (i.e., after depletion of the initially added $H_2O_2$) was inspired by Schwaiger et al (*25*). Residual activity was taken as the ratio of linear gradients fit to two segments of the $H_2O_2$ depletion data: (i) the first 10 s after reaction initiation, and (ii) a 10-s span following the second addition of $H_2O_2$. Ratios are reported as a value between 0 and 1. Mean residual activity was compared between variants by two-tailed unpaired *t* test.

*Stopped-flow redox kinetics*

The difference in fluorescence between the LPMO-$Cu^{2+}$ and LPMO-$Cu^{+}$ forms (*19*) was used to measure the kinetics of reduction by ascorbate and oxidation by $H_2O_2$, as previously described (*28*). All the experiments were carried out with a stopped-flow rapid spectrophotometer (SFM4000, BioLogic Science Instruments, Seyssinet-Pariset, France) coupled to a photomultiplier tube with an applied voltage of 600 V for detection. The excitation wavelength was set to 280 nm, and the fluorescence increase (for reduction) or decay (for oxidation) was collected with a 340 nm bandpass filter. All experiments were carried out at 25 ºC in 50 mM sodium phosphate, pH 7.0. To monitor reduction of $Cu^{2+}$ to $Cu^{+}$, LPMO-$Cu^{2+}$ (5 µM final concentration) was mixed with different concentrations of ascorbate (ranging from 5 to 250 µM final concentrations). For oxidation, double-mixing experiments were performed in two steps. In a first step, the LPMO-$Cu^{2+}$ (10 µM initial concentration) was mixed with one molar equivalent of L-cysteine for 10 seconds to form LPMO-$Cu^{+}$. In a second step, the *in situ* generated LPMO-$Cu^{+}$ was mixed with different concentrations of $H_2O_2$ (ranging from 25 to 800 µM after mixing) to monitor the decay of fluorescence. All stock solutions were deoxygenated by $N_2$ sparging using a Schlenk line and the working dilutions were subsequently prepared in sealed syringes in an anaerobic chamber. The stopped-flow rapid spectrophotometer was flushed with a large excess of deoxygenated buffer before coupling the sealed syringes and performing the experiments. BioKine32 software (V4.74.2, BioLogic) was used to fit the data to the single hyperbola $y = at + b + c \cdot e^{-k_{obs} \cdot t}$. Plots of $k_{obs}$ vs [ascorbate] or [$H_2O_2$] were fitted using linear least squares regression to obtain the apparent second order rate constant of the reduction or oxidation step ($k_{app}^{asc}$ or $k_{app}^{H2O2}$, respectively, assuming pseudo-first order conditions) with SigmaPlot v14.0.

*Stopped-flow UV-Vis detection of amino acid radicals*

All the experiments were carried out with a stopped-flow rapid spectrophotometer (SFM4000, BioLogic) coupled to a TIDAS® S 500 MCS UV/NIR 1910 (J&M Analytik AG, Essingen, Germany) diode array for detection of UV-Vis traces. Double mixing experiments were used to generate UV-Vis traces after mixing LPMO-$Cu^{+}$ with $H_2O_2$. To do this, LPMO-$Cu^{2+}$ (300 µM solution) was mixed with 1 molar equivalent of ascorbate, followed by ageing the reaction for 10 seconds in a delay line to ensure formation of LPMO-$Cu^{+}$. After that, the $Cu^{+}$ protein was mixed with 20 molar equivalents of $H_2O_2$, and UV-Vis traces were collected with 1.5 ms sampling between spectra to capture fast formation of radicals, and 75 ms sampling between spectra to capture long absorbance signal decay times. The instrument dead time was approximately 2.9 ms. All experiments were carried out at 4 ºC in 50 mM sodium phosphate, pH 7.0. All reagent solutions and the stopped-flow spectrophotometer were deoxygenated as described above. The $H_2O_2$ concentration of stock solutions was determined by measuring absorbance at 240 nm and using an extinction coefficient of 43.6 $M^{-1}$ $cm^{-1}$. Non-linear modelling of decay in absorbance at 415 nm was performed using a single exponential model of the form $y = a + b \cdot e^{-k_{obs} \cdot t}$. $k_{obs}$ was reported for Fig. 3J and Fig. 4E.

*Molecular dynamics simulations*

Inputs for the MD simulations and free-energy calculations were obtained from PDB entry 2BEM (*Sm*AA10A, chain C) or modelled using AlphaFold2 (*29*) for the ancestral LPMOs. The $Na^+$ ion modelled in the Cu-binding site in the 2BEM crystal structure was replaced by $Cu^{2+}$. This copper atom was also incorporated in the binding sites of the ancestral proteins. The models were prepared by manual editing and the pdb4amber program in the AmberTools23 package (*30*). The protonation state of each titratable amino acid side chain was predicted at pH 7.0 using the H++ (v3.2) software package (*31*), and the input PDB files were updated accordingly. Disulfide bridges were identified for Cys residues C14-C22 and C118-C135 in *Sm*AA10A, Anc3 and Anc4, for C14-C22, C49-C163 and C118-C135 in Anc2, and for C14-C22 and C49-C163 in Anc1. The program tleap (AmberTools23) was used to solvate the proteins in TIP3 water molecules using a box size that ensured 14 Å of solvent around the enzymes, generating input files for the MD-simulations. Chloride and sodium ions were added to the models to ensure charge neutrality and a 0.1 M salt concentration during the simulations.

For all the simulations, the AMBER ff14SBforce field was used for the protein (*32*) and the Joung/Cheatham ion parameters for TIP3P (*33*) were used for water and ions. The force-field parameters for $Cu^{2+}$ were adopted from previous work (*34*).

*Initial MD simulations.* For all models, the first stage involved a 5000-step energy minimization performed on the entire system. During the minimization, the non-hydrogen atoms of the enzymes were positionally restrained with a harmonic potential of 10 kcal $mol^{-1}$ $Å^{-2}$ for the first 2500 steps, then 1 kcal $mol^{-1}$ $Å^{-2}$ for the final 2500 steps. Restraints were only applied during the energy minimization procedure. Then, the systems were heated linearly from 0 to 300 K for 40 ps at constant volume using the Langevin thermostat with a collision frequency of 1 $ps^{-1}$. Density equilibrations were run at 300 K for 0.5 ns at a constant pressure of 1 atm using the Berendsen barostat with a pressure relaxation time of 1 ps. The final 100-ns equilibration step was carried out in the canonical (NVT) ensemble, where the number of particles, volume and temperature are held constant, using the weak coupling algorithm and a time constant of 10 ps to regulate the temperature. In all simulations, we used 2-fs time steps, periodic boundary conditions with a 12-Å cutoff for non-bonded interactions, and PME treatment of long-range electrostatics (*35*), whereas hydrogen atoms were constrained by the SHAKE algorithm

(*36*). Simulations were carried out using the CUDA version of PEMMD included in AMBER22 (*37*). Analysis of the resulting trajectories was performed using the cpptraj module included in AmberTools23 (*38*).

*Gaussian accelerated MD simulations.* To explore a wider conformational space in a more efficient way than conventional MD simulations, we performed Gaussian accelerated MD (GaMD) simulations, an enhanced sampling technique that flattens the potential energy surface of the system (*39*). Models of *Sm*AA10A or the ancestral LPMOs, with His133 in the HIE (histidine side chain protonated only at ε-nitrogen) or HID (histidine side chain protonated only at δ-nitrogen) protonation state, were equilibrated for 100 ns (as described above) before being subjected to GaMD simulations. The simulations were performed in the NVT ensemble with the parameters described above and run for 1 ms each, with the following GaMD-specific simulation flags: igamd = 3 (dual boost on both dihedral and total potential energy), iE = 1 (threshold energy set to the lower bound), ntcmdprep = $8\times10^5$ (number of preparation biasing MD simulation steps), nteb = $4\times10^6$ (number of biasing MD simulation steps), ntave = $2\times10^5$ (number of simulation steps used to calculate the average and standard deviation of potential energies), sigma0P = 6.0 (upper limit of the standard deviation of the first potential boost), sigma0D = 6.0 (upper limit of the standard deviation of the second potential boost). Coordinates were recorded every 1000 steps, resulting in trajectories with $5\times10^5$ frames for each simulation. The trajectories were analyzed using cpptraj (*38*), the values for the dihedral angles $\chi_1$ and $\chi_2$ were extracted for each frame, and the data were reweighted using the program PyReweighting (github.com/MiaoLab20/pyreweighting) by cumulant expansion to the second order (*40*, *41*).

*Constant-pH simulations.* The protonation state of His133 in *Sm*AA10A and the ancestral LPMOs was examined by constant-pH MD simulations (*42*). The total equilibration time before starting the constant-pH simulations was 100 ns. All explicit-solvent constant-pH simulations were carried out at 0.1 M salt concentration, using 100 MD steps between each protonation attempt and 200 MD steps of solvent relaxation following each attempt. For each model, we ran 16 parallel 200-ns replica exchange simulations at pH 1.5-9.0 (0.5-unit intervals), and replica exchange was attempted every 1000 steps. The data were processed with the program cphstats from AmberTools23. Eq. 1 was applied when fitting the Henderson-Hasselbach equation to the data ($f_d$ is the fraction of the protonated HIP state – the histidine side chain protonated at both ε- and δ-nitrogens).

(Equation 1) $$fd = \frac{1}{(1+10^{(n\cdot(pKa-pH))})}$$

*AlphaFold3 and Rosetta Relax*

Ancestral LPMO structures including copper ions were predicted using AlphaFold3 (*43*) with default parameters. For each prediction, the best-ranking structure according to the LDDT score was selected for analysis. Rosetta Relax (v2021.16) was used to refine backbone and sidechain conformations of the predicted structures to improve local geometry and resolve clashes (*44*). Input models were subjected to ten cycles of relaxation using the default FastRelax protocol with the ref2015 scoring function, with additional constraints imposed by the -in:auto_setup_metals flag. Five models were generated in each case, and the model with the lowest energy score was selected for analysis.

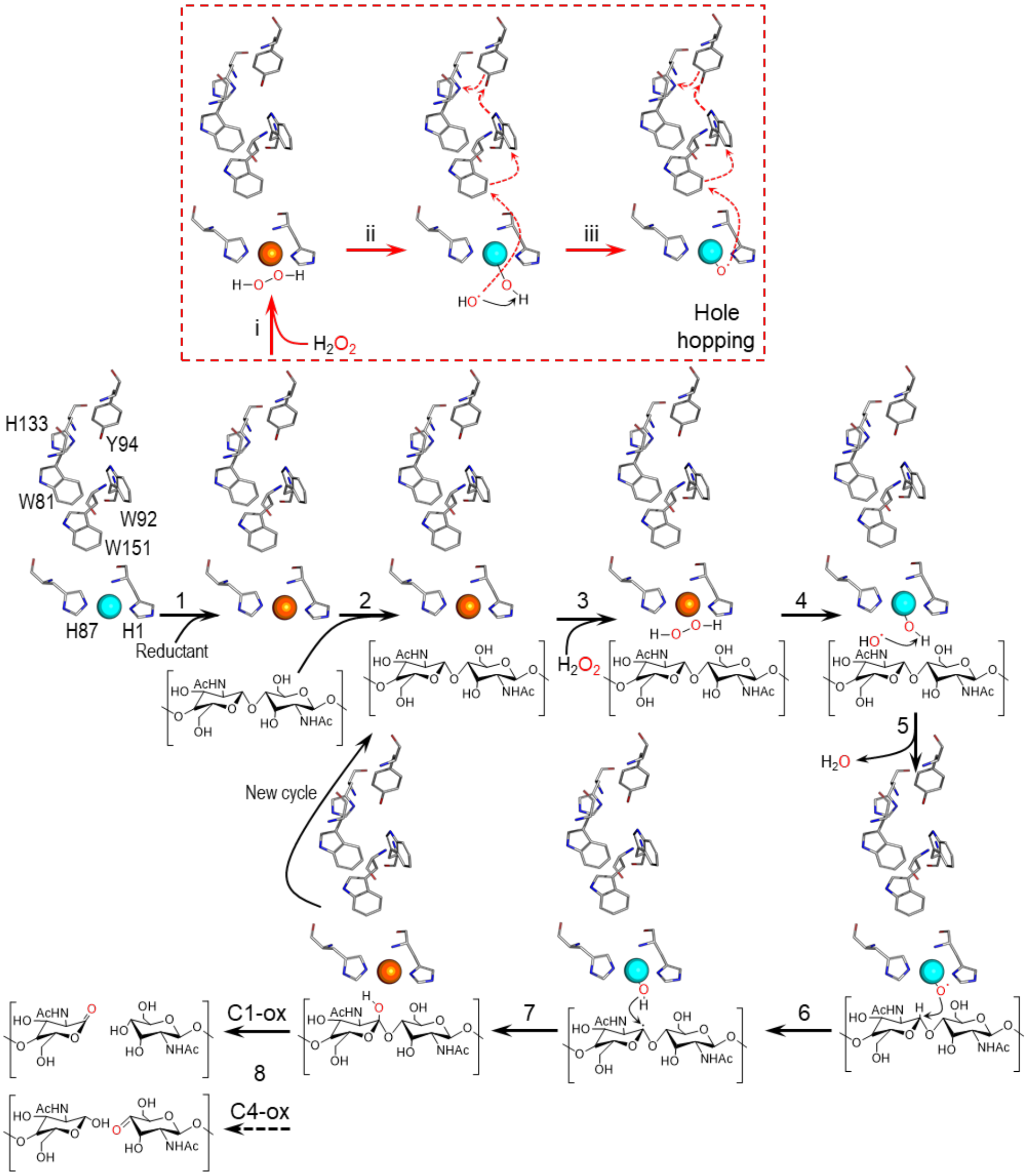

**Figure S1. Putative mechanism for the LPMO reaction.** The LPMO with a $Cu^{2+}$ cofactor (blue sphere) is reduced by a reductant (1) to generate the active $Cu^{+}$ form (orange sphere). The binding of the substrate (2) and $H_2O_2$ (3) initiates the productive oxidation mechanism, indicated by black arrows. This mechanism includes homolytic cleavage of $H_2O_2$ (4), formation of a copper-oxyl species that does the hydrogen abstraction (5,6) and a "rebound" step leading to substrate hydroxylation (7). The mechanism shown is compatible with mechanistic studies carried out by multiple research groups (*28*, *45–47*). The hydroxylated glycosidic bond is unstable and rearrangement will lead to chain cleavage (*48*), generating either a lactone, for C1-oxidizing LPMOs, such as the LPMOs studied here, or a 4-keto sugar (for C4 oxidizing LPMOs). These oxidized products are in equilibrium with their hydrated forms, an aldonic acid or a 4-gemdiol, respectively. In the absence of a bound substrate, the reaction of the reduced LPMO with $H_2O_2$ can lead to a futile hole hopping reaction (red box, **i** to **iii**) involving the aromatic core of the LPMO (red arrows). This hole hopping mechanism can be initiated by the hydroxyl radical generated after homolytic cleavage of $H_2O_2$ (**ii**) or by subsequently emerging radical species such as the Cu-oxyl (**iii**). The transference of these holes is key to avoid damage in the histidine brace and to protect the LPMO from autocatalytic inactivation (*26*). The red arrows illustrate one possible route for hole dissipation; additional routes are possible (*26*).

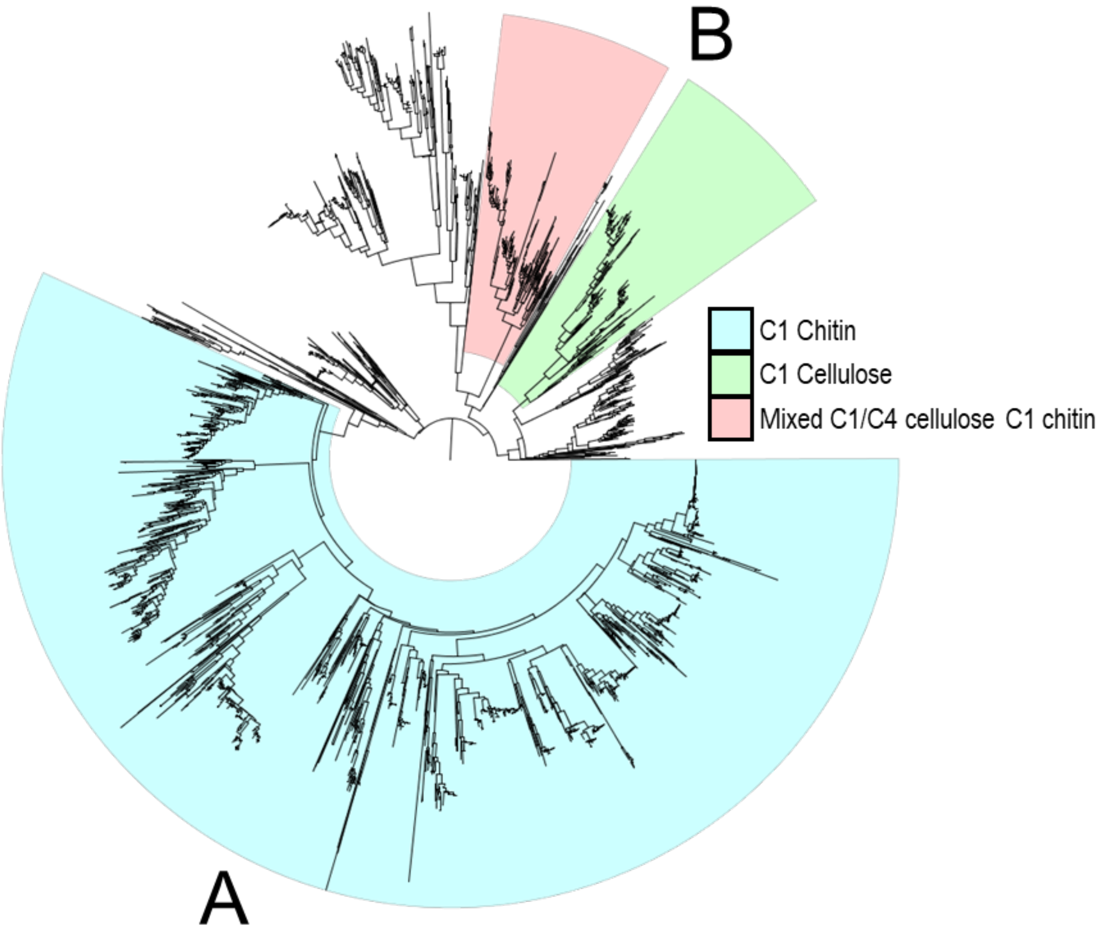

**Figure S2. Phylogenetic tree of 3332 AA10 LPMO sequences extracted from dbCAN** (*1*). Signal peptides and CBMs were removed, and the catalytic domains were aligned using MAFFT with automatic selection of parameters. The tree was built using fasttree, with default parameters. The topology of the tree for the reduced subset of AA10 LPMOs (main text Fig. 1A) is consistent with previous studies (*3*), showing primarily two large clades (named A and B). Clade A corresponds to strictly chitin-active LPMOs, while clade B is a mixture of strictly chitin-active and strictly cellulose-active LPMOs and LPMOs with a mixed chitin and cellulose activity. In this study we focused on clade A, which contains chitin-active LPMOs from Bacilli, Gammaproteobacteria and Actinomycetes, covering a wide taxonomic variety of organisms with medical and ecological relevance. Cellulose-active and mixed activity AA10 LPMOs have so far only been described in Actinomycetes. This phylogenetic tree was used to select the 159 sequences for the ancestral sequence reconstruction while maintaining the phylogenetic diversity found in the sequence space of AA10 LPMOs. 25 amino acid sequences of characterized LPMOs (*2*, *49*) were used to identify clades with known substrate specificities.

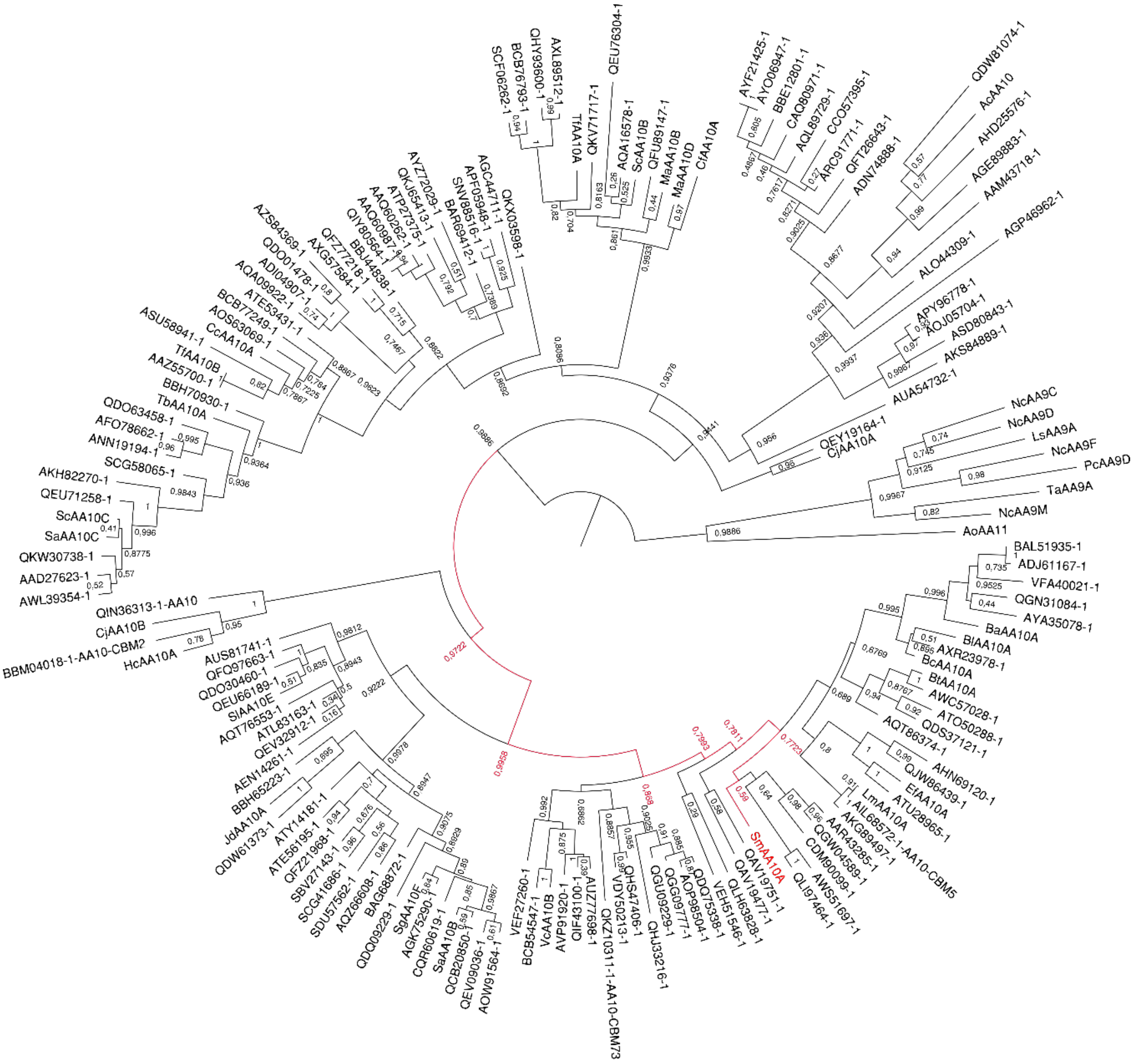

**Figure S3. Phylogenetic tree of AA10 LPMOs including accession numbers and bootstrap support values**. A complete phylogenetic tree corresponding to the reduced schematic shown in Fig. 1A. 159 sequences were used to build the tree, using 7 AA9 LPMOs and 1 AA11 LPMO as root. The sequences were aligned using MAFFT, and the tree was built with PhyML using the parameters for the best model calculated with Prottest3.4. Here, the GenBank accession numbers of all LPMOs and the bootstrap values for each node (100 independent replicas) are included. Note that the majority of the nodes have bootstrap values > 0.75, indicating well supported clades. For a list of all accession numbers see Table S10.

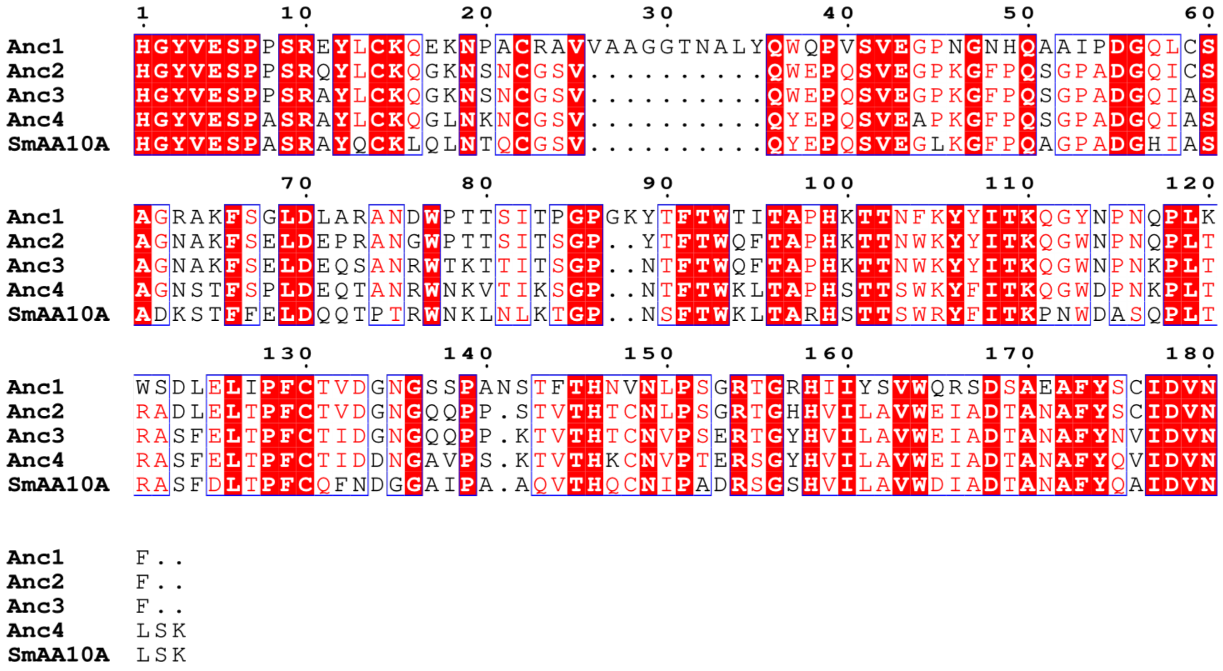

**Figure S4. Sequence analysis of selected ancestral LPMOs and *Sm*AA10A.** The multiple sequence alignment was performed with MUSCLE as implemented in MEGA X (*50*) and colored to show patterns of conserved amino acids using ESPrit 3 (*51*) (https://espript.ibcp.fr). The sequences used were as follows:

Anc1:
HGYVESPPSREYLCKQEKNPACRAVVAAGGTNALYQWQPVSVEGPNGNHQAAIPDGQLCSA
GRAKFSGLDLARANDWPTTSITPGPGKYTFTWTITAPHKTTNFKYYITKQGYNPNQPLKWSD
LELIPFCTVDGNGSSPANSTFTHNVNLPSGRTGRHIIYSVWQRSDSAEAFYSCIDVNF

Anc2:
HGYVESPPSRQYLCKQGKNSNCGSVQWEPQSVEGPKGFPQSGPADGQICSAGNAKFSELDEP
RANGWPTTSITSGPYTFTWQFTAPHKTTNWKYYITKQGWNPNQPLTRADLELTPFCTVDGNG
QQPPSTVTHTCNLPSGRTGHHVILAVWEIADTANAFYSCIDVNF

Anc3:
HGYVESPPSRAYLCKQGKNSNCGSVQWEPQSVEGPKGFPQSGPADGQIASAGNAKFSELDEQ
SANRWTKTTITSGPNTFTWQFTAPHKTTNWKYYITKQGWNPNKPLTRASFELTPFCTIDGNGQ
QPPKTVTHTCNVPSERTGYHVILAVWEIADTANAFYNVIDVNF

Anc4:
HGYVESPASRAYLCKQGLNKNCGSVQYEPQSVEAPKGFPQSGPADGQIASAGNSTFSPLDEQT
ANRWNKVTIKSGPNTFTWKLTAPHSTTSWKYFITKQGWDPNKPLTRASFELTPFCTIDDNGAV
PSKTVTHKCNVPTERSGYHVILAVWEIADTANAFYQVIDVNLSK

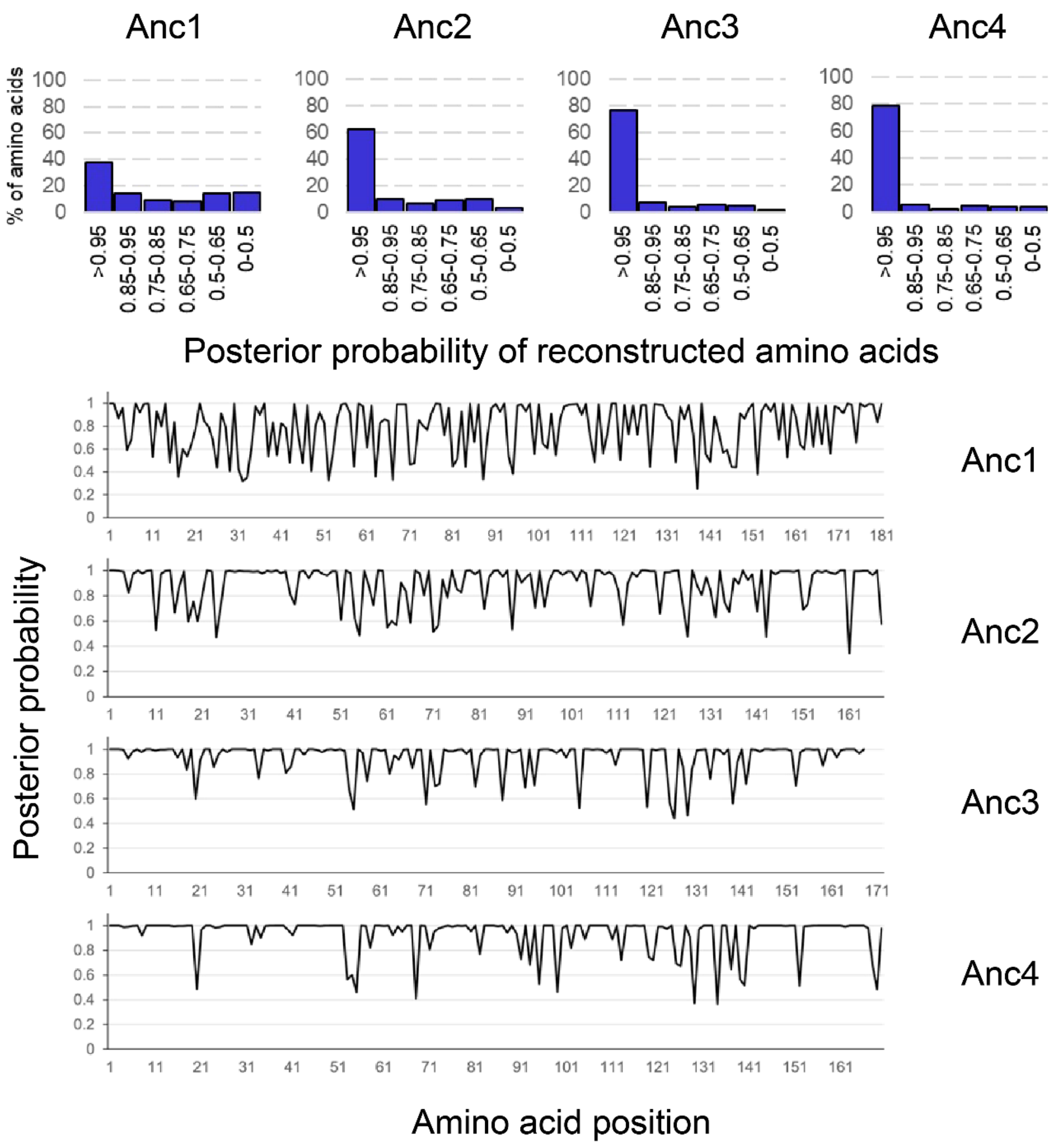

**Figure S5. Posterior probability analysis for the most probable sequence reconstructed at each selected node.** Bar plots on top represent the fraction (%) of amino acids within a given posterior probability range. Bottom panels represent the posterior probability for each position in the reconstructed sequences. The amino acid with the highest posterior probability was selected for each position.

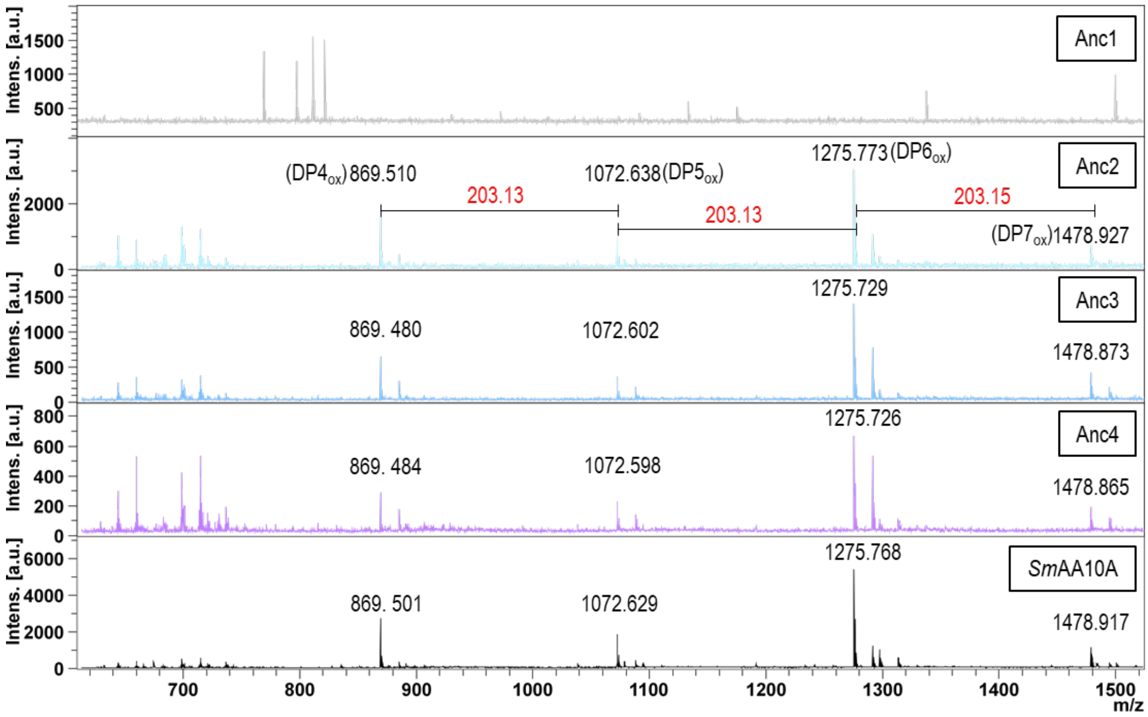

**Figure S6. MALDI-TOF MS analysis of chitin oxidation by ancestral enzymes and *Sm*AA10A.** The figure shows signals corresponding to soluble native and oxidized LPMO products in the DP4 to DP7 range (DP, degree of polymerization). The products were observed as sodium or potassium adducts of the lactone or, mostly, the aldonic acid form, or as sodium adducts of sodium salts of the aldonic acid form, where the latter is diagnostic for C1 oxidation. The signals labeled with their *m/z* value correspond to the sodium adduct of the aldonic acid. For example, 1072 is the sodium adduct of the aldonic acid form of C1-oxidized chitopentaose. The masses of the products differ by 203 Da per *N*-acetylglucosamine unit. The reactions were run for 6 hours in 50 mM sodium phosphate, pH 7.0, containing 10 g $L^{-1}$ chitin, 1 mM ascorbate and 1 µM LPMO, at 37 °C. Substrate solubilization did not occur in control reactions containing LPMO but lacking the reductant.

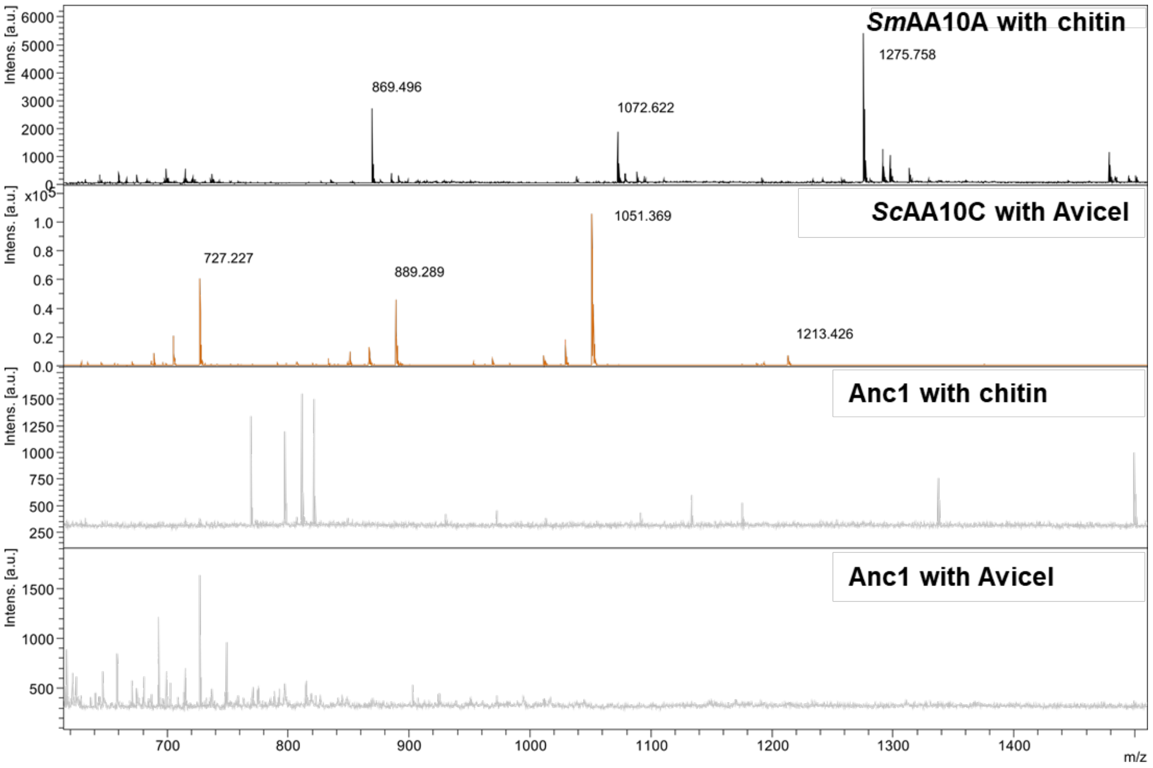

**Figure S7. MALDI-TOF MS analysis of chitin or cellulose oxidation by ancestral Anc1 and *Sm*AA10A or *Sc*AA10C.** *Sm*AA10A and *Sc*AA10C (*14*) were used as a positive controls for chitin and cellulose (Avicel) oxidation, respectively. Major signals for oxidized chitooligosaccharides (DP4-DP7) or cello-oligosaccharides (DP4-DP7) are marked by their *m/z* value. For *Sm*AA10A the signals labeled with their *m/z* value correspond to the sodium adduct of the aldonic acid. For *Sc*AA10C (cellulose), the sodium adduct of the sodium salt of the aldonic acid dominates and this signal is labelled. The reactions were run for 6 hours in 50 mM sodium phosphate, pH 7.0, containing 10 g $L^{-1}$ chitin or 10 g $L^{-1}$ Avicel, 1 mM ascorbate and 1 μM LPMO, at 37 °C.

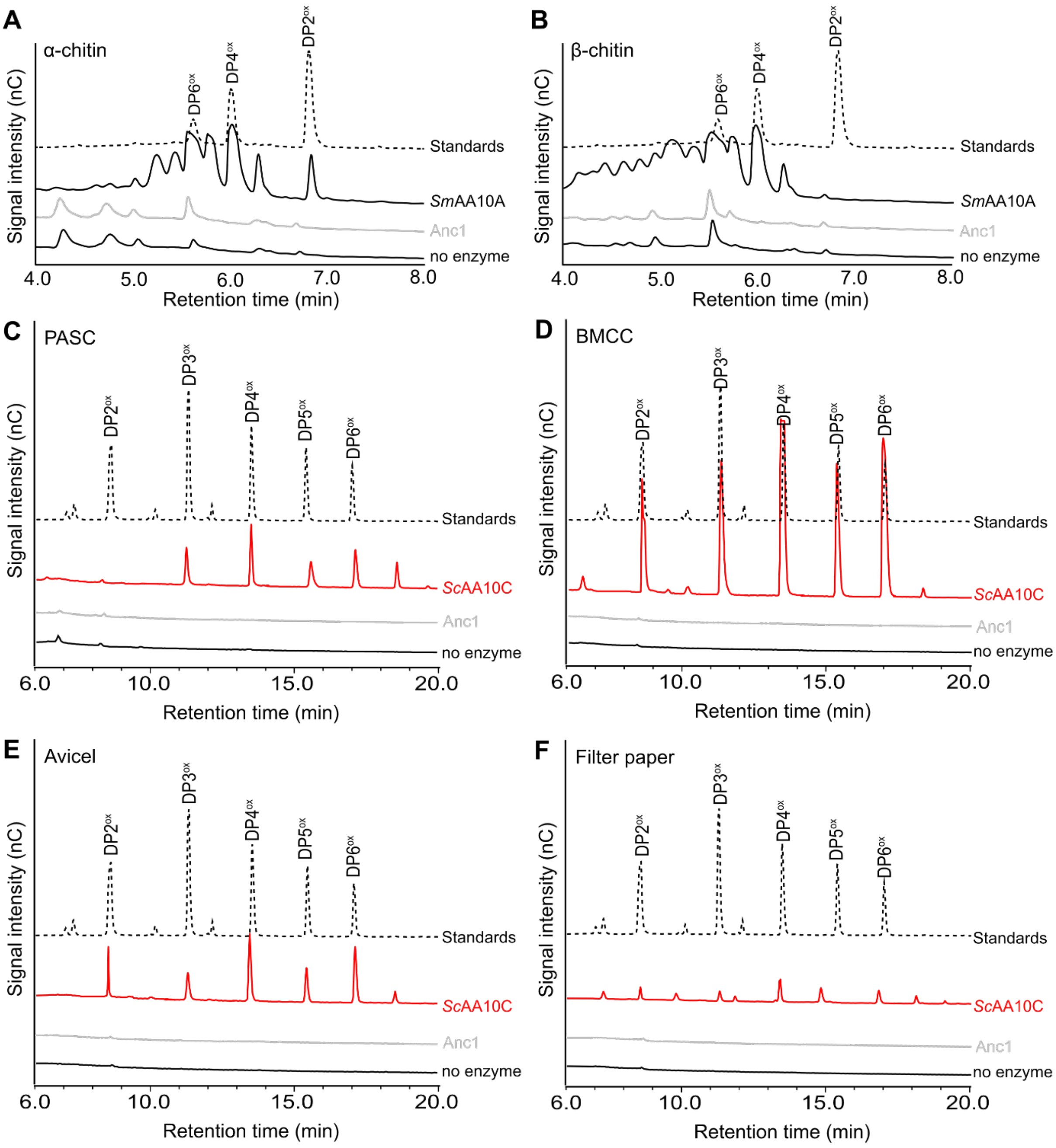

**Figure S8. HPAEC-PAD chromatograms of oxidized products released from chitinous and cellulosic substrates using Anc1.** Reactions were carried out at 30 ºC in 50 mM Tris, pH 8.0, containing 5% glycerol, using 1 µM LPMO, 1 mM ascorbate and 0.5% (w/v) α- or β-chitin, 0.2% (w/v) bacterial microcrystalline cellulose (BMCC), 0.5% (w/v) filter paper (Whatman no.1), 1% (w/v) Avicel® PH-101, or 0.2% (w/v) phosphoric acid swollen cellulose (PASC). Samples were taken after 48 hours of reaction.

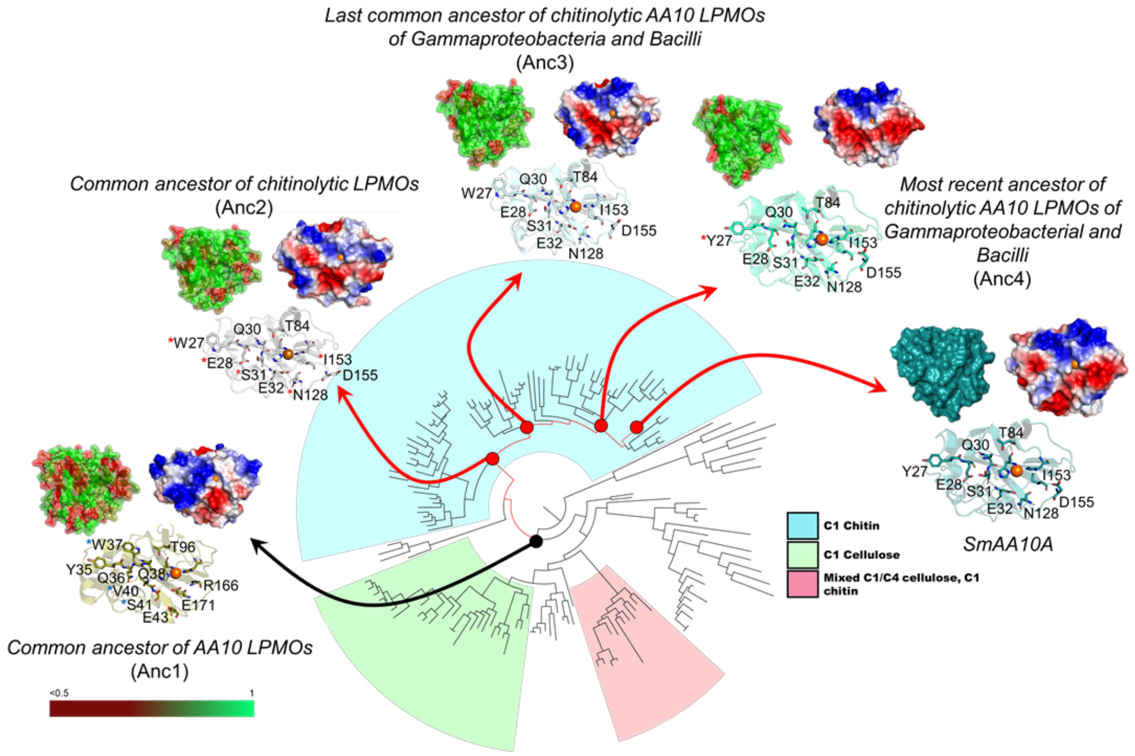

**Figure S9. Surface analysis of ancestral AA10s**. The figure depicts the phylogenetic tree of Figure 1 of the main manuscript with the selected nodes of the chitin-active lineage (red arrows) and the common ancestor, Anc1 (black arrow). For each node, the posterior probability of the ancestral reconstruction (red-green scale, lower left corner), and the surface charge (red - negative, blue - positive) are shown. Table S2 provides posterior probabilities for selected surface residues. Blue asterisks indicate amino acids that are lost in evolution or rearranged due to deletions. Red asterisks indicate amino acids that appear in evolution.

**Supplementary discussion I - Evolution of the substrate-binding surface**

The analysis of the binding surface shown in Fig. S9 highlights that the surface changed both in terms of amino acid composition and overall charge in the chitin-active lineage. While it is not possible to link the evolution of charge to chitin-binding, since this has not been studied, variations at the amino acid level include one notable change, namely a transition from Trp27 present in Anc2 and Anc3 to Tyr27 in Anc4 and *Sm*AA10A (Table S2). The importance of this residue for substrate binding is well documented by mutational studies (*52*, *53*). It has been shown that mutation of this tyrosine in *Sm*AA10A reduces binding to chitin (*52*). Nakagawa and colleagues (*53*) demonstrated that mutation of this residue from Tyr to Trp in a small chitin-active LPMO from *Streptomyces griseus* has an effect on how the LPMO synergizes with chitinases, but they were not able to demonstrate significant effects of the mutation on binding to α-chitin and β-chitin.

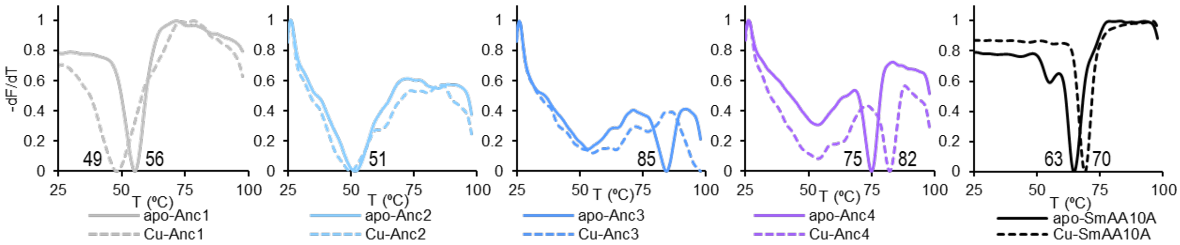

**Figure S10. Melting temperatures of ancestral LPMOs and *Sm*AA10A with and without $Cu^{2+}$ bound.** The SyproOrange® thermal shift assay was used to determine the apparent $T_m$, which is the temperature at which *dF/dT* reaches its maximum. Reactions were carried out in 50 mM Tris, pH 7.0, using 30 µM LPMO, in quadruplicates. Apparent $T_m$ values are shown in the figure (averages of 4 replicates).

**Supplementary discussion II – Limiting factors in LPMO reactions**

LPMO reactions may be set up in two principally different manners, with or without an external source of $H_2O_2$. In both reactions a reductant is needed to reduce and, thus, activate the LPMO. In reactions without an external source of $H_2O_2$, i.e., conditions that are commonly used in the field, the LPMO reaction is reductant driven. Under these conditions, sometimes referred to as "apparent monooxygenase" or "*in situ* $H_2O_2$ producing" conditions, the reaction is limited by *in situ* production of $H_2O_2$ through oxidation of the reductant, which happens both abiotically (sometimes referred to as "auto-oxidation") and as a result of the oxidase activity of the LPMO (*54*). In the case of *Sm*AA10, which has very low oxidase activity (*55*), and at the pH used here with ascorbate (*56*), $H_2O_2$ availability is primarily determined by abiotic oxidation of the reductant.

Reactions with externally added $H_2O_2$, will normally run much faster, depending on the rate of $H_2O_2$ addition and will not lead to stoichiometric consumption of reductant (*57*, *58*). Reduced LPMOs that meet $H_2O_2$ in the absence of substrate can catalyze a futile peroxidase reaction which may damage the enzyme and lead to enzyme inactivation (*57*). The chance of an enzyme being damaged depends on its tendency to engage in this off-pathway reaction and the chance that such reaction leads to oxidative damage. These tendencies and chances vary among LPMOs (*27*). Because of this, LPMO reactions, either reductant-driven or fuelled by externally added $H_2O_2$, may lead to enzyme inactivation to an extent that depends on the substrate concentration and the enzyme's affinity for the substrate.

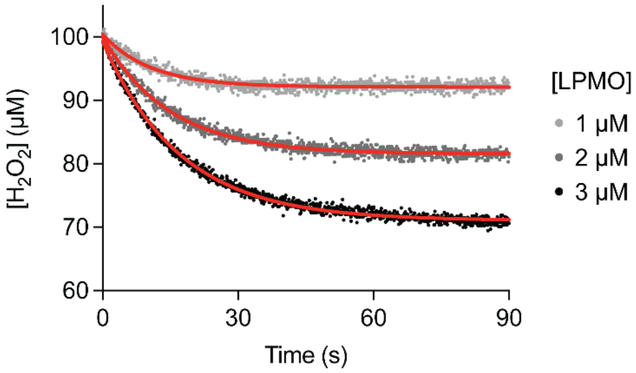

**Figure S11. $H_2O_2$ consumption in the absence of chitin.** The graph shows typical progress curves for $H_2O_2$ consumption by *Sm*AA10A in the absence of chitin underlying the data presented in Fig. 3, C-D and Fig. 4, F-G of the main manuscript. Only single experiments are shown for the sake of clarity. Each curve was fit using non-linear regression, from which values for the extent and rate of $H_2O_2$ consumption were extracted as described in the Methods. $R^2 > 0.9$ for all curves. Control reactions without ascorbate or without LPMO showed no detectable $H_2O_2$ consumption over 2 min (not shown). Reactions were monitored using a rotating disk electrode with an angular velocity of 50 $s^{-1}$, at 30 °C in 50 mM potassium phosphate, pH 7.0, 100 mM KCl and 5 µM EDTA as detailed in the Methods. Data points were recorded at 0.08 s intervals. EDTA (at low concentration) was added in these reactions to scavenge free copper released from damaged LPMO molecules.

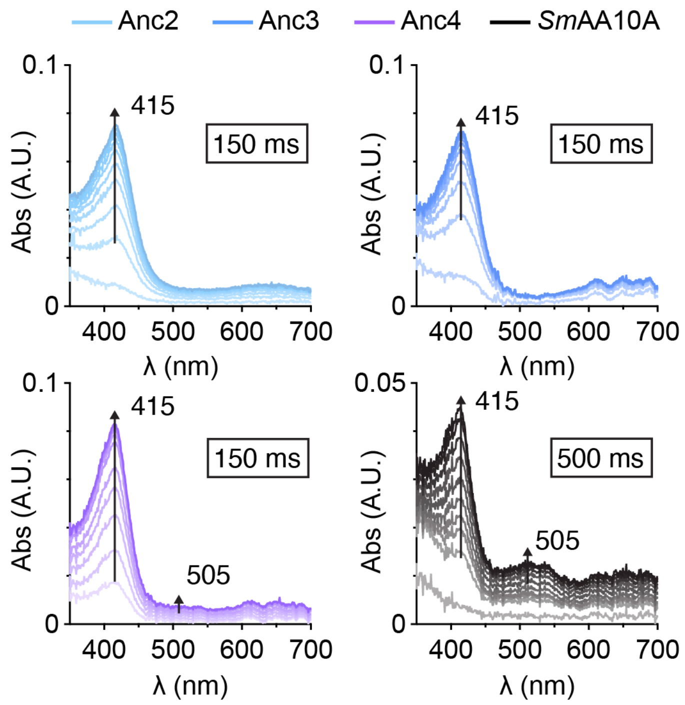

**Figure S12. UV-vis absorbance spectra showing amino acid radical formation for Anc2-4 and *Sm*AA10A.** The graphs show absorbance spectra for Anc2, Anc3, Anc4 and *Sm*AA10A following reactions between LPMO-$Cu^+$ and $H_2O_2$ in the absence of chitin. All reactions contained 75 µM LPMO, that was first reduced by mixing with 1 molar equivalent of ascorbate, then with 20 molar equivalents of $H_2O_2$ in 50 mM sodium phosphate, pH 7.0, at 4 °C. Data are means ($N \geq 3$) from the same experiments used to produce Fig. 3, E-H; error bars are not shown for clarity. Ten spectra are shown for each enzyme, spanning from 20 ms to the time of maximum absorbance at 415 nm. The time to maximum absorbance is shown in the box in each panel. Note the different scale of the ordinate axis for *Sm*AA10A.

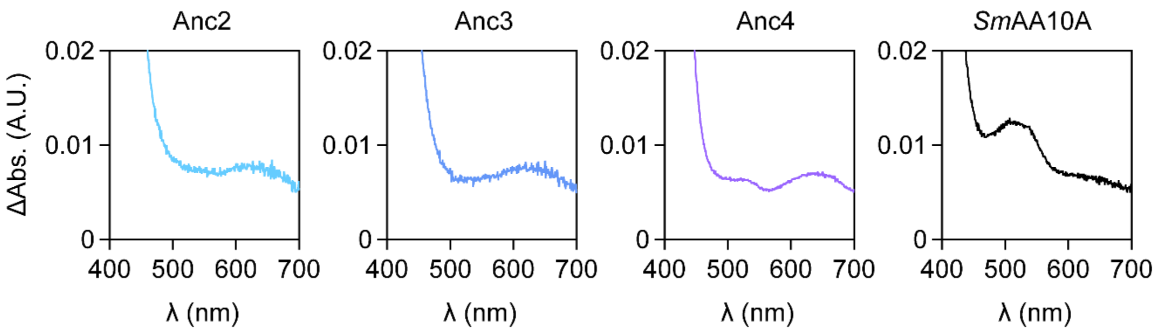

**Figure S13. Spectral subtractions of absorbance at 500 ms minus 20 ms after mixing LPMO-Cu$^+$ with $H_2O_2$.** The graphs are derived from average spectra (N ≥ 3 technical replicates) derived from the same datasets as those presented in Fig. 3E-H of the main text. Zooms of the ordinate axis are shown to focus on spectral features around 500 nm. The broad peak around 640 nm likely corresponds to a change in absorbance caused by oxidation of the LPMO from the Cu$^+$ to the Cu$^{2+}$ state as expected (*21*), while the broad peak with maximum at ~505 nm in Anc4 and SmAA10A likely corresponds to a change in absorbance caused by formation of a Trp$^•$ (see main text). All reactions contained 75 µM LPMO that was first reduced by mixing with 1 molar equivalent of ascorbate, then mixed with 20 molar equivalents of $H_2O_2$ in 50 mM sodium phosphate, pH 7.0, at 4 °C.

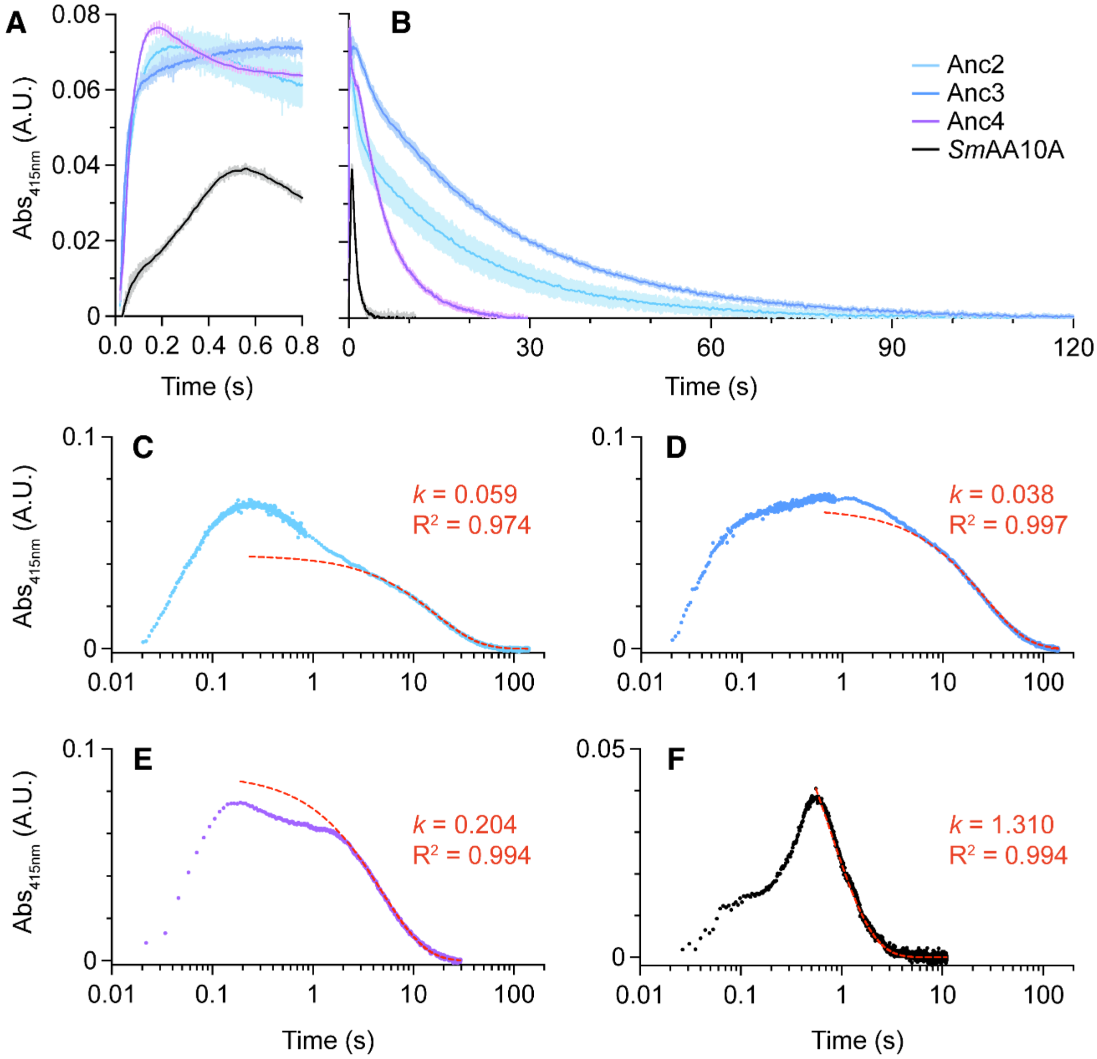

**Figure S14. Time-resolved traces and non-linear modelling for the Tyr˙ feature observed in ancestral LPMOs and *Sm*AA10A.** The plots on the top row represent (**A**) the first 800 ms, to focus on formation, or (**B**) the full runs to show full decay of the Tyr˙ feature (415 nm) shown in Fig. 3 of the main text. All reactions contained 75 µM LPMO that was first reduced by mixing with 1 molar equivalent of ascorbate then mixed with 20 molar equivalents of $H_2O_2$ in 50 mM sodium phosphate, pH 7.0, at 4 °C. The shadowed areas represent SDs ($N \geq 3$ technical replicates). Panels **C-F** show single example runs for Anc2-4 and *Sm*AA10A overlaid with a single exponential model as detailed in the Methods (red dashed lines). For each enzyme, the signal decay from the time of maximum absorbance showed a complex shape suggesting multiple underlying processes. However, the final phase of each decay fit closely to a single exponential decay model. Rates ($k$; $s^{-1}$) and $R^2$ values are indicated in each case. Rates extracted from each model are reported in Fig. 3J of the main text. The colour coding indicated in (B) applies to all panels.

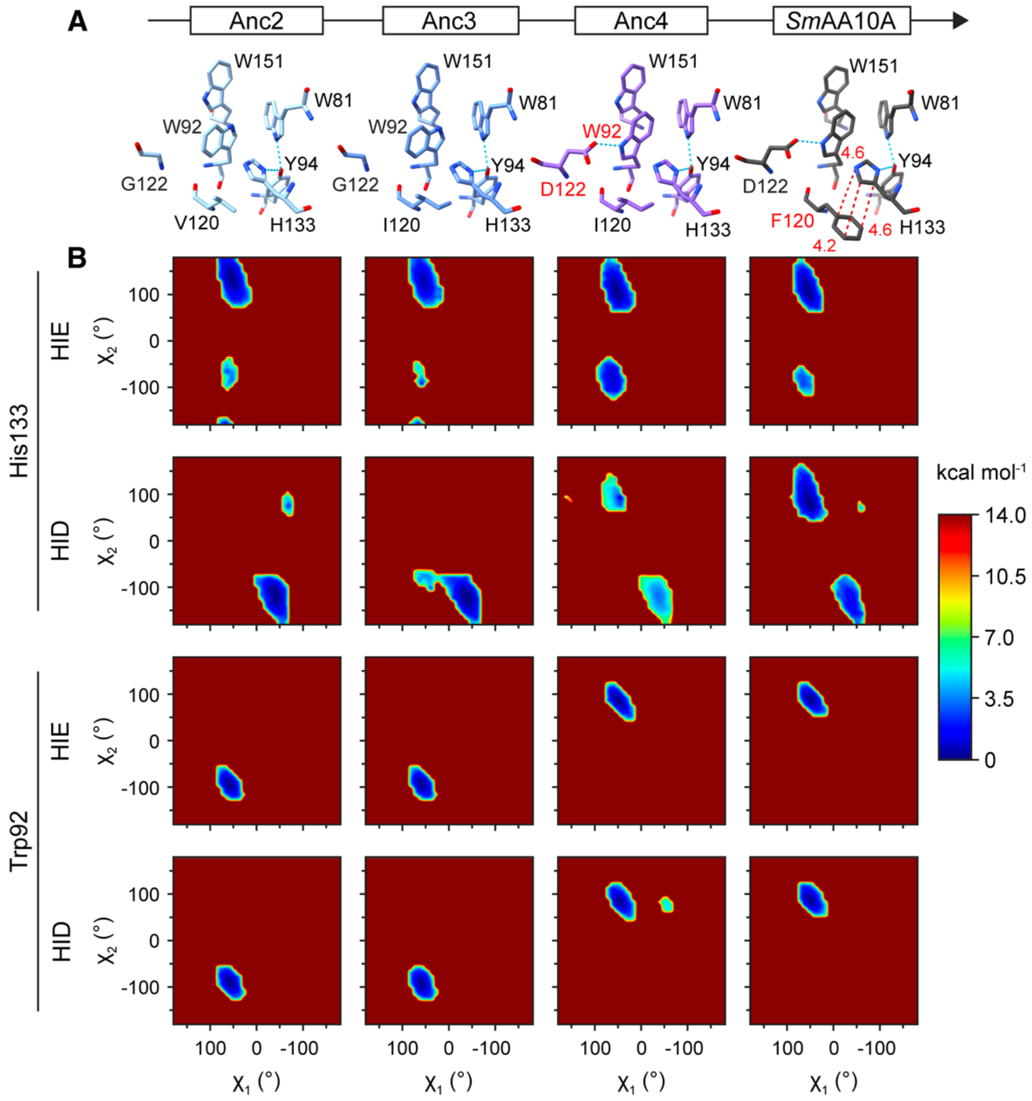

**Figure S15. Energetics of Trp92 and His133 conformations in Anc2-4 and *Sm*AA10A probed by Gaussian accelerated MD simulations.** (**A**) Predicted models of Anc2-4 and SmAA10A showing residues thought to be involved in hole hopping (Trp151, Trp92, Trp81, Tyr94, His133) and residues at positions 120 and 122, the identity of which change over evolution. Residue labels in red mark the point in evolution in which they first appear or are first predicted to occupy the orientation found in SmAA10A. Distances (red) are in angstroms. (**B**) The potential energy surfaces as a function of the side-chain dihedral angles $\chi_1$ and $\chi_2$ (see Fig. S16) for His133 in the HIE and HID protonation states, and for Trp92 with His133 in HIE or HID in Anc2-4 and *Sm*AA10A. The free energy profile (potential of mean force) is shown in kcal mol$^{-1}$ according to the indicated coloring scheme. The deep red color indicates high-energy states that are not likely to be populated, whereas blue color indicates energetically favourable states. The graphs show that in the first two steps in evolution, each protonation state of His133 fluctuates around one conformational state, with dihedral angles, $\chi_1$ and $\chi_2$, centered at -60º, +125º for HIE and +50º, -125º for HID. When Asp122 appears in Anc4 and *Sm*AA10A, His133 adopts two conformational states, centered around -60º, +125º and -50º, -110º for HIE, and -60º, +100º and +50º, -125º for HID. The conformation of Trp92 clearly changes during evolution: going from Anc3 to Anc4, when Asp122 appears, there is a clear transition from the ancestral state (-60º, -100º) to the "flipped" state (-50º, +75º). These conformational states are independent of the His133 protonation state.

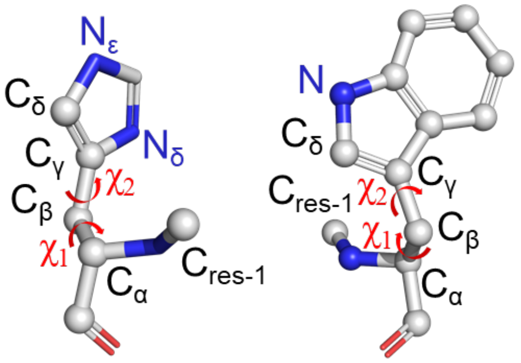

**Figure S16. Dihedral angles ($\chi$1 and $\chi$2) for H133 and W92.** Angles analyzed in the Gaussian accelerated MD simulations depicted in Fig. S15. Residue numbering applies to Anc2-4 and *Sm*AA10A.

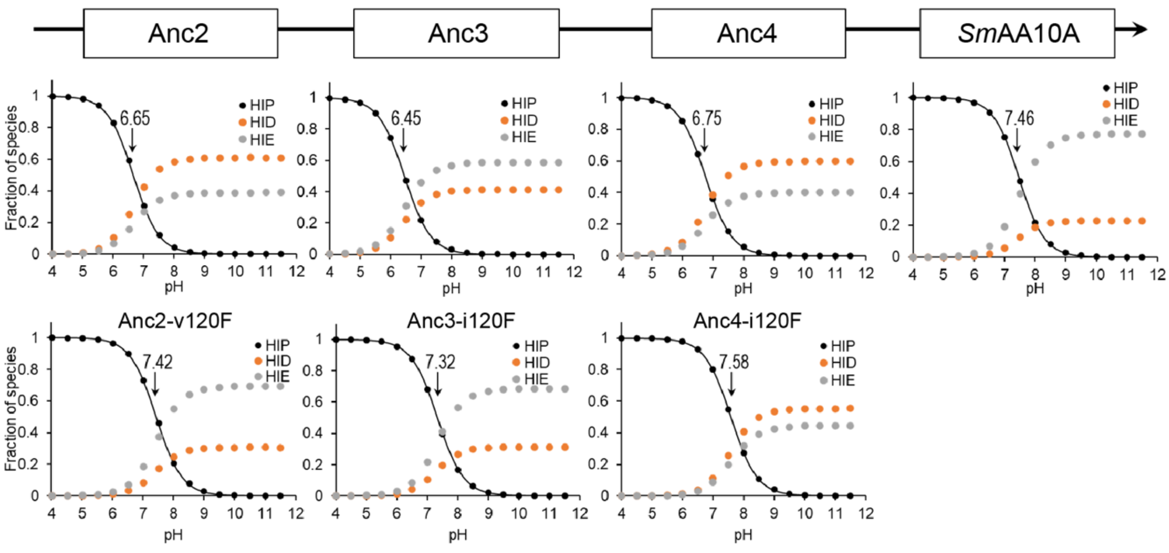

**Figure S17. Estimation of titration midpoints and protonation states for His133.** Constant-pH MD simulations of Anc2-4, *Sm*AA10A and the Anc2_v120F, Anc3_i120F and Anc4_i120F variants. The data points show the fractions of HIP (black), HIE (gray) and HID (orange). The data for HIP were fitted to the Henderson-Hasselbach equation (solid line) using Eq. 1 (see Methods). Black arrows indicate estimated p$K_a$ values. His133 has an apparent p$K_a$ of 6.65 and a slight preference for protonation of the Nδ in Anc2, while in *Sm*AA10A, the apparent p$K_a$ is 7.46, and protonation of Nε is clearly preferred. In between, the picture remains comparable to Anc2 (Fig. S17), indicating that the appearance of Phe120, in *Sm*AA10A, was key for the observed change in the properties of His133. We also simulated variants Anc2-v120F, Anc3-i120F and Anc4-i120F, confirming that the presence of Phe120 increases the apparent p$K_a$ of His133 by almost one unit and makes the Nε protonation state of the imidazole ring more favorable.

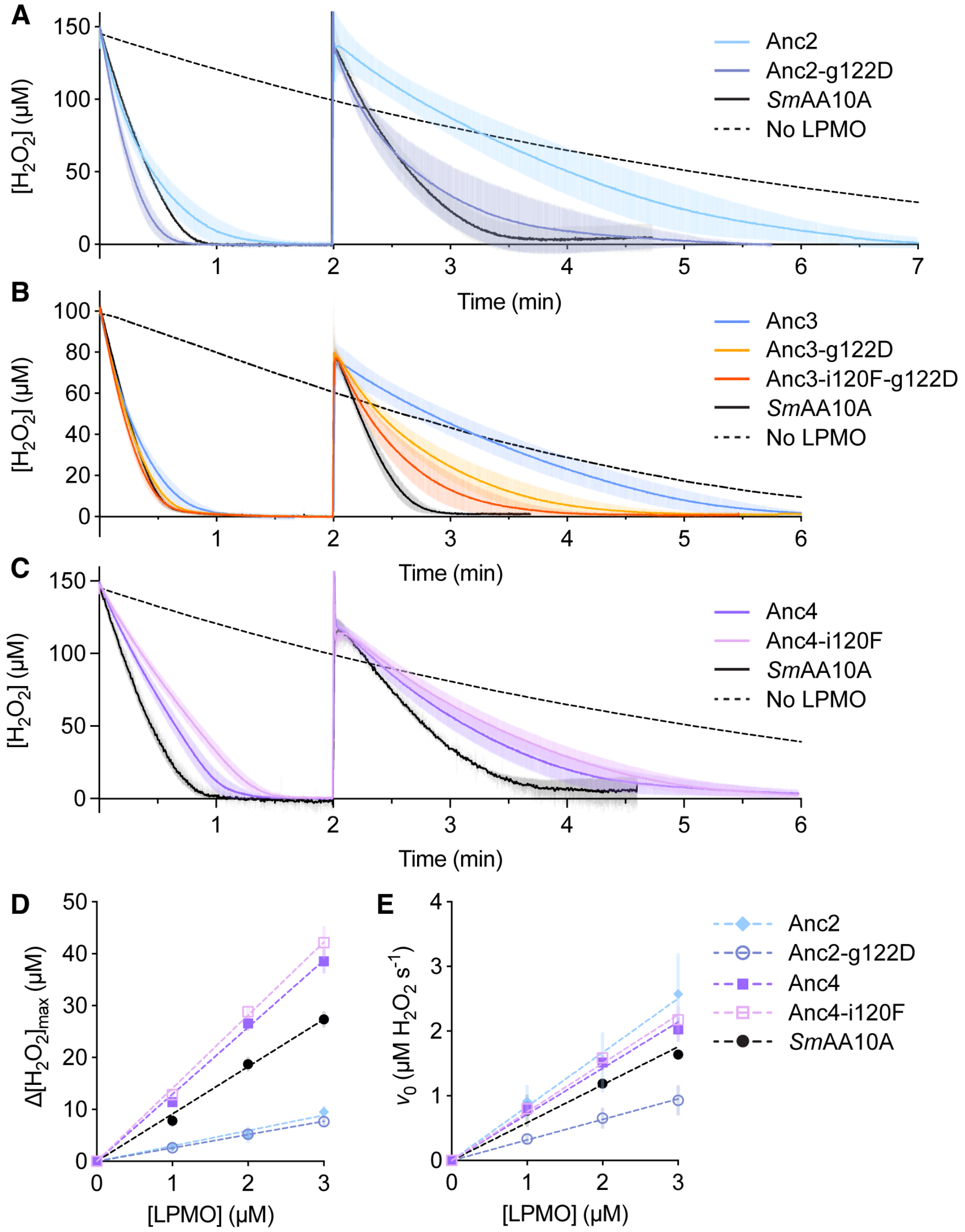

**Figure S18. Impact of introducing Phe120 and/or Asp122 in Anc2-4 on reactions with added $H_2O_2$.** Real-time $H_2O_2$ consumption for Anc2 (**A**), Anc3 (**B**) and Anc4 (**C**) in the presence of chitin, including mutants with changes at positions 120 and/or 122 discussed in the main text. Curves shown in (B) are identical to those shown in Fig. 4B of the main text. All reactions contained 1 µM LPMO, 10 g $L^{-1}$ chitin, 1 mM ascorbate and 100 mM KCl in 50 mM potassium phosphate, pH 7.0, at 37 °C. The initial concentration of $H_2O_2$ was 150 µM in (A) and (C), but 100 µM in (B). A second addition of $H_2O_2$ was made after depletion of the initially added $H_2O_2$. Data are means ($N \geq 3$ independent experiments); shaded areas indicate SDs. (**D**) Maximum $H_2O_2$ turned over in reactions at 30 °C with 100 mM KCl, 5 µM EDTA, 200 µM ascorbate and 100 µM initial $H_2O_2$ in the absence of chitin. Error bars show SDs ($N \geq 2$ independent experiments). Dashed lines show linear regression of the data; slopes define $n_{max}$ (listed in Table S8). (**E**) Initial rate of $H_2O_2$ turnover derived from the experiments performed to produce (D). Dashed lines show linear regression of the data; slopes represent the apparent turnover number (TN, $s^{-1}$) (Table S8). The legend in panel E also applies to panel D. We also attempted to produce the Anc2-v120F-g122D double mutant but were unable to express this protein.

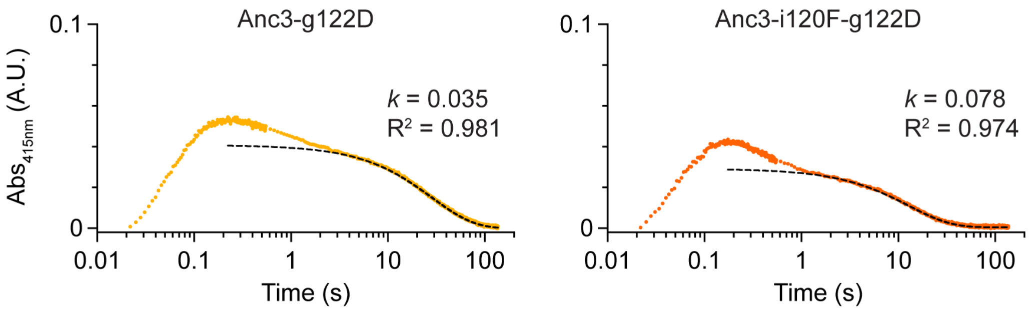

**Figure S19. Time-resolved traces and non-linear modelling for the $Tyr^{\bullet}$ feature observed in mutants of Anc3.** Plots show single example runs for the mutants Anc3-g122D and Anc3-i120F-g122D overlaid with a single exponential model as detailed in the Methods (black dashed lines). Reactions contained 75 µM LPMO that was first reduced by mixing with 1 molar equivalent of ascorbate then mixed with 20 molar equivalents of $H_2O_2$. Reactions were performed in 50 mM sodium phosphate, pH 7.0, at 4 °C in the absence of chitin. For each enzyme, the signal decay from the time of maximum absorbance showed a complex shape suggesting multiple underlying processes. However, the final phase of each decay fit closely to a single exponential decay model. Rates ($k$; $s^{-1}$) and $R^2$ values are indicated in each case. Rates extracted from each model are reported in Fig. 4E of the main text.

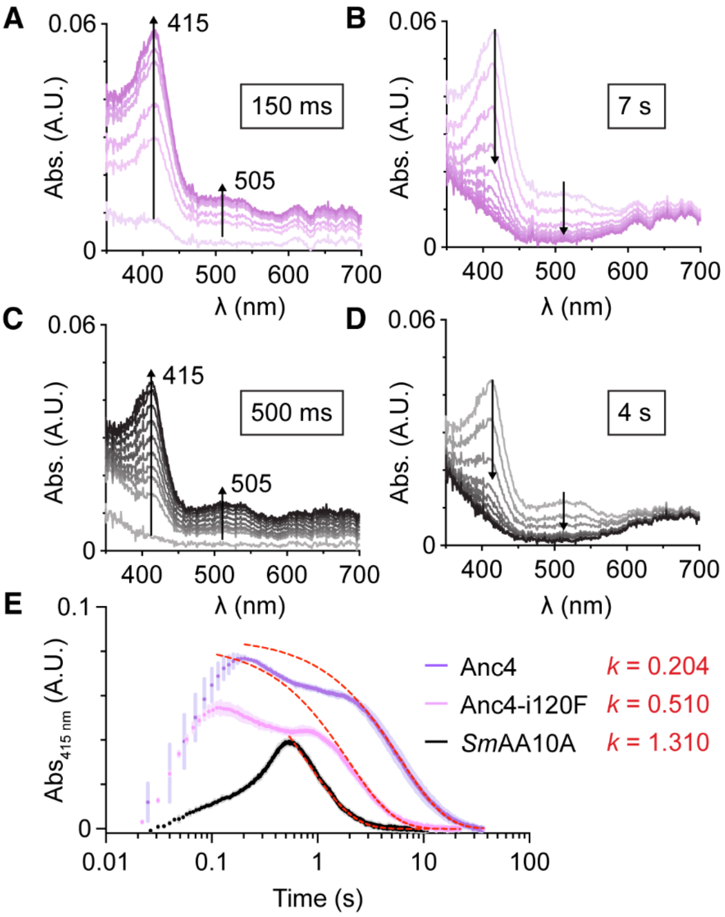

**Figure S20. UV-vis based detection of amino acid radicals in Anc4-i120F.** Panels (**A**) and (**B**) show formation (A) and decay (B) absorbance spectra for the Anc4-i120F variant. Panels (**C**) and (**D**) show formation (C) and decay (D) spectra for SmAA10A, reproduced here to illustrate the similarities to Anc4-i120F. The data shown in (C) is the same as that shown in Fig. S12D and the data shown in (D) is the same as that shown in Fig. 3H. Traces in (A-D) are means ($N \geq 4$ technical replicates); standard deviations are not shown for the sake of clarity. Panel (**E**) shows time-resolved absorbance at 415 nm derived from the data for Anc4-i120F and *Sm*AA10A in panels (A-D), supplemented with data for Anc4 (formation data in Fig. S12C and decay data in Fig. 3G). Traces are means ± SDs ($N \geq 4$ technical replicates). Signal decay at 415 nm was fit to a single exponential decay model (red dashed lines) starting from the time of maximum signal intensity as described in the Methods. The rates ($k$; $s^{-1}$) are indicated in each case. All reactions contained 75 µM LPMO that was first reduced by mixing with 1 molar equivalent of ascorbate then mixed with 20 molar equivalents of $H_2O_2$ in 50 mM sodium phosphate, pH 7.0, at 4 °C.

**Table S1. Posterior probabilities for the amino acids in the His-brace.** Residue numbers correspond to *Sm*AA10A. Please note that in most previous publications, residue numbering in *Sm*AA10A included the signal peptide, which means that residue numbers were 27 higher (for example, the N-terminal His1 was referred to as His28).

| | His-brace | |
|---|---|---|
| Anc1 | 1.000 | 1.000 |
| Anc2 | 1.000 | 1.000 |
| Anc3 | 1.000 | 1.000 |
| Anc4 | 1.000 | 1.000 |
| *Sm*AA10A | H1 | H87 |

**Table S2. Posterior probabilities for amino acids important for chitin binding.** Residues were identified as important for binding to chitin according to (*34*, *59*). Residue numbers correspond to *Sm*AA10A. Chosen amino acids in ambiguous positions are highlighted in bold. Please note that in most previous publications, residue numbering in *Sm*AA10A included the signal peptide, which means that residue numbers were 27 higher (for example, the N-terminal His1 was referred to as His28).

| | Surface | | | |
|---|---|---|---|---|
| Anc1 | W:0.999 | **Q:0.536**<br>E:0.139<br>M:0145<br>N:0.084 | **V:0.545**<br>I:0.299<br>L:0.101 | T:0.980 |
| Anc2 | W:0.999 | 0.992 | Q:0.995 | T:0.999 |
| Anc3 | W:1.000 | E:1.000 | Q:1.000 | T:1.000 |
| Anc4 | Y:1.000 | E:1.000 | Q:1.000 | T:1.000 |
| *Sm*AA10A | Y27 | E28 | Q30 | T84 |

**Table S3. Posterior probabilities for amino acids participating in the main hole hopping route identified in SmAA10A** (*26*). Residue numbers correspond to *Sm*AA10A. Chosen amino acids in ambiguous positions are highlighted in bold. Please note that in most previous publications, residue numbering in *Sm*AA10A included the signal peptide, which means that residue numbers were 27 higher (for example, the N-terminal His1 was referred to as His28).

| | Hole hopping route | | | | |
|---|---|---|---|---|---|
| Anc1 | F:0.997 | W:0.999 | **F:0.910**<br>W:0.059<br>Y:0.027 | **Y:0.867**<br>F:0.130 | **H:0.442**<br>M:0.350<br>I:0.074 |
| Anc2 | F:1.000 | W:1.000 | **W:0.977**<br>F:0.022 | Y:0.998 | H:0.998 |
| Anc3 | F:1.000 | W:1.000 | W:1.000 | Y:1.000 | H:1.000 |
| Anc4 | F:1.000 | W:1.000 | W:1.000 | Y:1.000 | H:1.000 |
| *Sm*AA10A | F160 | W151 | W92 | Y94 | H133 |

**Table S4. Posterior probabilities for amino acids defining the 2nd coordination sphere of the copper.** Residue numbers correspond to *Sm*AA10A. Chosen amino acids in ambiguous positions are highlighted in bold. Please note that in most previous publications, residue numbering in *Sm*AA10A included the signal peptide, which means that residue numbers were 27 higher (for example, the N-terminal His1 was referred to as His28).

| | 2nd coordination sphere | | | | |
|---|---|---|---|---|---|
| Anc1 | **E:0.483**<br>A:0.100<br>Q:0.084 | **R:0.965**<br>K:0.026 | **D:0.979**<br>N:0.012 | **E:0.958**<br>D:0.056 | F:0.997 |
| Anc2 | E:0.996 | **I:0.731**<br>V:0.264 | D:0.999 | **N:0.982**<br>D:0.012 | F:1.000 |
| Anc3 | E:1.000 | **I:0.707**<br>V:0.293 | D:1.000 | N:1.000 | F:1.000 |
| Anc4 | E:1.000 | I:0.995 | D:1.000 | N:1.000 | F:1.000 |
| *Sm*AA10A | E33 | I153 | D155 | N158 | F160 |

**Table S5. Observed rate constants for the production of $H_2O_2$ and redox potentials (mV vs NHE) of the ancestral LPMOs and *Sm*AA10A.** The production of $H_2O_2$ was measured using the Amplex Red assay (*21*) as described in the Methods. Rates were corrected for the rate in reactions without LPMO. Redox potentials were measured using the TMP assay (*60*), in 50 mM Tris, pH 8.0, using 35 µM LPMO and 150 µM TMP. Reported values are the means ± SDs ($N = 3$ independent measurements).

| | Observed rate constant ($s^{-1}$) | Redox potential (mV) |
|---|---|---|
| Anc2 | 0.0018 ± 0.0003 | 244 ± 7 |
| Anc3 | 0.0115 ± 0.0041 | 251 ± 6 |
| Anc4 | 0.0122 ± 0.0016 | 237 ± 3 |
| *Sm*AA10A | 0.0014 ± 0.0011 | 244 ± 7 |

**Table S6. Thermal inactivation of Anc3 and *Sm*AA10A.** A 2 µM solution of *Sm*AA10A or Anc3 in 50 mM sodium phosphate, pH 7.0, was incubated for 10 mins at either 75 ºC or 90 ºC, and, after cooling down and centrifugation, LPMO activity was assessed by adding 1 mM 2,6-DMP and 100 µM $H_2O_2$ (final concentrations). The numbers represent the activity as $\Delta Abs_{473nm}$ $min^{-1}$, with the error as the standard deviation of three independent experiments. The percentages represent the residual activity after temperature incubation, using as 100% their activity prior to incubation. N.D.: not detected, implying complete loss of activity.

| Temperature | 75 ºC | 90 ºC |
| --- | --- | --- |
| *Sm*AA10A | 0.0005 ± 0.0002 (1%) | N.D. |
| Anc3 | 0.0109 ± 0.0003 (61%) | 0.0046 ± 0.0001 (30%) |

**Table S7. Summary of reduction and reoxidation rates presented in Fig. 3.** The table shows the apparent second-order rate constants for reduction of LPMO-$Cu^{2+}$ by ascorbate, presented in Fig. 3A, and oxidation of LPMO-$Cu^{+}$ by $H_2O_2$, presented in Fig. 3B, for Anc2-4 and *Sm*AA10A. Data are based on pseudo-first order rate constants for the change in fluorescence intensity plotted against the concentration of ascorbate or $H_2O_2$, respectively. The ratio of the second-order rate constants is also reported. All reported rates are the average of at least three technical replicates with standard deviations shown.

| | $k_{app}^{asc}$ ($M^{-1}$ $s^{-1}$) | $k_{app}^{H2O2}$ ($M^{-1}$ $s^{-1}$) | Ratio red/ox |
|---|---|---|---|
| Anc2 | 210000 ± 370 | 13900 ± 10 | 15 |
| Anc3 | 145000 ± 100 | 23300 ± 20 | 6.2 |
| Anc4 | 238000 ± 720 | 9200 ± 10 | 26 |
| *Sm*AA10A | 381000 ± 570 | 6900 ± 30 | 55 |

**Table S8. Summary of $n_{max}$ values and turnover numbers for the ascorbate-peroxidase reaction.** The table shows kinetic parameters for the ascorbate-peroxidase reaction presented in Figs. 3 and 4 and Fig. S18, which were determined by real-time monitoring of $H_2O_2$ consumption. Example curves are shown in Fig. S11, and these curves were fitted to a single exponential model as detailed in the Methods. The average number of peroxidase reactions that one LPMO can catalyze before inactivation, $n_{max}$, was estimated as the maximum $H_2O_2$ turned over in the reaction ($\Delta[H_2O_2]_{max}$) divided by the enzyme concentration (*27*). The apparent turnover number (TN, $s^{-1}$) was estimated from the initial rate of $H_2O_2$ consumption ($v_0$, µM $s^{-1}$) divided by the enzyme concentration (*27*). Background $H_2O_2$ consumption (in the absence of LPMO) could not be detected on the experimental timescale (two minutes) so was not used to correct $v_0$. All reported values are means ($N \geq 2$); errors are SEM calculated using the LINEST function in Excel. Reactions contained 0, 1, 2 or 3 µM LPMO, 100 µM initial $H_2O_2$, 100 mM KCl and 5 µM EDTA in 50 mM potassium phosphate, pH 7.0, at 30 °C. Reactions were initiated with 200 µM ascorbate.

| | $n_{max}$ | TN ($s^{-1}$) |
|---|---|---|
| Anc2 | 3.27 ± 0.10 | 0.95 ± 0.06 |
| Anc3 | 3.14 ± 0.15 | 1.33 ± 0.09 |
| Anc4 | 12.91 ± 0.23 | 0.71 ± 0.02 |
| *Sm*AA10A | 9.12 ± 0.21 | 0.59 ± 0.03 |
| Anc2-g122D | 2.57 ± 0.08 | 0.32 ± 0.02 |
| Anc3-g122D | 3.35 ± 0.11 | 0.80 ± 0.05 |
| Anc3-i120F-g122D | 3.35 ± 0.14 | 0.76 ± 0.05 |
| Anc4-i120F | 14.07 ± 0.27 | 0.75 ± 0.02 |

**Table S9. Posterior probabilities for amino acids interacting with the main hole hopping route.** Residue numbers correspond to *Sm*AA10A. Chosen amino acids in ambiguous positions are highlighted in bold. Please note that in most previous publications, residue numbering in *Sm*AA10A included the signal peptide, which means that residue numbers were 27 higher (for example, the N-terminal His1 was referred to as His28).

| | | |
|---|---|---|
| Anc1 | **V: 0.846**<br>I: 0.146<br>F: 0 | G: 0.970<br>D: 0.009 |
| Anc2 | **V: 0.662**<br>I: 0.321<br>F: 0.002 | G: 0.988<br>D: 0.005 |
| Anc3 | V: 0.467<br>**I: 0.533**<br>F: 0 | G: 0.996<br>D: 0.002 |
| Anc4 | V: 0.249<br>**I: 0.724**<br>F: 0.025 | G: 0.002<br>D: 0.992 |
| *Sm*AA10A | F120 | D122 |

**Table S10. LPMO accession numbers.** List of NCBI's Gene database accession numbers of LPMOs used for the phylogenetic trees in Fig. 1 and Fig. S3. For LPMOs having a specific name, composed of a species name followed by AA10 and a letter indicating the isoform (i.e. *Sm*AA10A), the corresponding Gene accession number is given in the second column. If a Gene database accession number was not available, the UniProt accession number is provided and is labeled with *.

| Name in the tree | Gene/Uniprot |
|---|---|
| BAL51935.1 | |
| ADJ61167.1 | |
| VFA40021.1 | |
| QGN31084.1 | |
| AYA35078.1 | |
| *Ba*AA10A | CBI42985.1 |
| *Bl*AA10A | AAU39477.1 |
| AXR23978.1 | |
| *Bc*AA10A | AAP09751.1 |
| *Bt*AA10A | ACW83015.1 |
| AWC57028.1 | |
| ATO50288.1 | |
| QDS37121.1 | |
| AQT86374.1 | |
| AHN69120.1 | |
| QJW86439.1 | |
| *Ef*AA10A | AAO80225.1 |
| ATU28965.1 | |
| *Lm*AA10A | CAD00545.1 |
| AIL68572.1 | |
| AKG89497.1 | |
| AAR43285.1 | |
| QGW04589.1 | |
| CDM90099.1 | |
| AWS51697.1 | |
| QLI97464.1 | |
| *Sm*AA10A | AAU88202.1 |
| QAV19751.1 | |
| QAV19477.1 | |
| QLH63828.1 | |
| VEH51546.1 | |
| QDQ75338.1 | |
| AOP98504.1 | |
| QGG09777.1 | |
| QGU09229.1 | |
| QHJ33216.1 | |
| QHS47406.1 | |
| VDY50213.1 | |
| QKZ10311.1 | |
| AUZ77698.1 | |
| QIF43100.1 | |
| AVP91920.1 | |
| *Vc*AA10B | ABU72648.1 |
| BCB54547.1 | |
| VEF27260.1 | |
| AOW91564.1 | |
| QEV09036.1 | |
| QCB20850.1 | |
| *Sa*AA10B | CAJ89556.1 |
| CQR60619.1 | |
| AGK75290.1 | |
| *Sg*AA10F | BAG23684.1 |

| | |
|---|---|
| QDQ09229.1 | |
| BAG68872.1 | |
| AQZ66608.1 | |
| SDU57562.1 | |
| SCG41686.1 | |
| SBV27143.1 | |
| QFZ21968.1 | |
| ATE56195.1 | |
| ATY14181.1 | |
| QDW61373.1 | |
| *Jd*AA10A | ACV09037.1 |
| BBH65223.1 | |
| AEN14261.1 | |
| QEV32912.1 | |
| ATL83163.1 | |
| AQT76553.1 | |
| *Sl*AA10E | EOY47895.1 |
| QEU66189.1 | |
| QDO30460.1 | |
| QFQ97663.1 | |
| AUS81741.1 | |
| *Hc*AA10A | ABC27701.1 |
| BBM04018.1 | |
| *Cj*AA10B | ACE84760.1 |
| QIN36313.1 | |
| AWL39354.1 | |
| AAD27623.1 | |
| QKW30738.1 | |
| *Sa*AA10C | CAJ90160.1 |
| *Sc*AA10C | CAB61600.1 |
| QEU71258.1 | |
| AKH82270.1 | |
| SCG58065.1 | |
| ANN19194.1 | |
| AFO78662.1 | |
| QDO63458.1 | |
| *Tb*AA10A | D6Y7U3* |
| BBH70930.1 | |
| AAZ55700.1 | |
| *Tf*AA10B | AAZ55700.1 |
| ASU58941.1 | |
| *Cc*AA10A | AAF22274.1 |
| AOS63069.1 | |
| BCB77249.1 | |
| ATE53431.1 | |
| AQA09922.1 | |
| ADI04907.1 | |
| QDO01478.1 | |
| AZS84369.1 | |
| AXG57584.1 | |
| QFZ77218.1 | |
| BBJ44838.1 | |
| QIY80564.1 | |
| AAQ60987.1 | |
| AAQ60262.1 | |
| ATP27375.1 | |
| QKJ65413.1 | |
| AYZ72029.1 | |
| BAR69412.1 | |
| SNV88516.1 | |

| | |
|---|---|
| APF05948.1 | |
| AGC44711.1 | |
| QKX03598.1 | |
| SCF06262.1 | |
| BCB76793.1 | |
| QHY93600.1 | |
| AXL89512.1 | |
| *Tf*AA10A | AAZ55306.1 |
| QKV71717.1 | |
| QEU76304.1 | |
| AQA16578.1 | |
| *Sc*AA10B | CAB61160.1 |
| QFU89147.1 | |
| *Ma*AA10B | D9SZQ3* |
| *Ma*AA10D | D9T1F0* |
| *Cf*AA10A | ADG73094.1 |
| AYF21425.1 | |
| AYO06947.1 | |
| BBE12801.1 | |
| CAQ80971.1 | |
| AQL89729.1 | |
| CCO57395.1 | |
| ARC91771.1 | |
| QFT26643.1 | |
| ADN74888.1 | |
| QDW81074.1 | |
| *Ac*AA10 | O70709* |
| AHD25576.1 | |
| AGE89883.1 | |
| AAM43718.1 | |
| ALO44309.1 | |
| AGP46962.1 | |
| APY96778.1 | |
| AOJ05704.1 | |
| ASD80843.1 | |
| AKS84889.1 | |
| AUA54732.1 | |
| QEY19164.1 | |
| *Cj*AA10A | ACE83992.1 |
| *Nc*AA9C | EAA36362.1 |
| *Nc*AA9D | CAD21296.1 |
| *Ls*AA9A | ALN96977.1 |
| *Nc*AA9F | CAD70347.1 |
| *Pc*AA9D | BAL43430.1 |
| *Ta*AA9A | ACS05720.1 |
| *Nc*AA9M | EAA33178.1 |
| *Ao*AA11 | BAE61530.1 |
